# AcrIIA22 is a novel anti-CRISPR that impairs SpyCas9 activity by relieving DNA torsion of target plasmids

**DOI:** 10.1101/2020.09.28.317578

**Authors:** Kevin J. Forsberg, Danica T. Schmidtke, Rachel Werther, Deanna Hausman, Barry L. Stoddard, Brett K. Kaiser, Harmit S. Malik

## Abstract

To overcome CRISPR-Cas defense systems, many phages and mobile genetic elements encode CRISPR-Cas inhibitors called anti-CRISPRs (Acrs). Nearly all mechanistically characterized Acrs directly bind their cognate Cas protein to inactivate CRISPR immunity. Here, we describe AcrIIA22, an unconventional Acr found in hypervariable genomic regions of *Clostridial* bacteria and their prophages from the human gut microbiome. Uncovered in a functional metagenomic selection, AcrIIA22 does not bind strongly to SpyCas9 but nonetheless potently inhibits its activity against plasmids. To gain insight into its mechanism, we obtained an X-ray crystal structure of AcrIIA22, which revealed homology to PC4-like nucleic-acid binding proteins. This homology helped us deduce that *acrIIA22* encodes a DNA nickase that relieves torsional stress in supercoiled plasmids, rendering them less susceptible to SpyCas9, which is highly dependent on negative supercoils to form stable R-loops. Modifying DNA topology may provide an additional route to CRISPR-Cas resistance in phages and mobile genetic elements.

## Introduction

CRISPR-Cas systems in bacteria and archaea confer sequence-specific immunity against invading phages and other mobile genetic elements (MGEs)^1,2^. In response, MGEs can circumvent CRISPR-Cas systems by evading CRISPR immunity. In its simplest form, evasion requires only a single mutation to a CRISPR target site, which allows a phage or MGE to escape immune recognition^3^. However, CRISPR-Cas systems routinely acquire new spacer sequences to target new sites within phage and MGE genomes^1^. This means that any single-site evasion strategy is likely to be short-lived. Thus, phages also employ forms of CRISPR-Cas evasion that are less easily subverted. For instance, some jumbophages assemble a proteinaceous, nucleus-like compartment around their genomes upon infection, allowing them to overcome diverse bacterial defenses, including CRISPR-Cas and restriction-modification (RM) systems^4,5^. Similarly, other phages decorate their DNA genomes with diverse chemical modifications, which can prevent Cas nucleases from binding their target sequence, such as the glucosylated cytosines used by phage T4 of *Escherichia coli*^6^.

MGEs may also overcome CRISPR-Cas systems by inactivating, rather than evading, CRISPR immunity. MGEs encode diverse CRISPR-Cas inhibitors called anti-CRISPRs (Acrs), which allow them to overcome CRISPR-Cas systems and infect otherwise immune hosts^7^. Most known Acrs bind Cas proteins and inhibit Cas activity by either restricting access to target DNA, preventing necessary conformational changes, or inactivating critical CRISPR-Cas components^8,9^. The direct inactivation of Cas proteins by Acrs has proven an effective and widespread strategy for overcoming CRISPR immunity^10^.

Recent genetic, bioinformatic, and metagenomic strategies have identified many Acrs that independently target the same CRISPR-Cas system^7–10^. Yet, most CRISPR-Cas systems are not inhibited by known Acrs^10^. Thus, many undiscovered strategies to inhibit or evade CRISPR-Cas systems probably exist in nature. Indeed, over half of the genes in an average phage genome have no known function^11^. To uncover new counter-immune strategies, we recently devised a high-throughput functional metagenomic selection to find genes that protect a target plasmid from *Streptococcus pyogenes* Cas9 (SpyCas9), the variant used most frequently for genome editing^12^. Our selection strategy was designed to reveal any gene capable of overcoming SpyCas9 activity in this system, regardless of mechanism. With this approach, we previously described a new phage inhibitor of SpyCas9, called AcrIIA11, which acts via a novel mechanism and is prevalent across human gut microbiomes^12^.

Here, we describe AcrIIA22, which was the second most common Acr candidate recovered from our original functional selection. AcrIIA22 encodes a 54 amino acid protein that impairs SpyCas9 activity. We observe that homologs of *acrIIA22* are found in hypervariable loci in phage and bacterial genomes. Unlike most other Acrs, AcrIIA22 does not bind strongly to SpyCas9 *in vitro*. Instead, guided by an X-ray crystal structure of AcrIIA22, we show that AcrIIA22 encodes a DNA nickase. By nicking a supercoiled plasmid substrate and relieving its torsional stress, AcrIIA22 renders the target less susceptible to SpyCas9 activity. AcrIIA22 thus represents a novel mechanism of SpyCas9 evasion, which capitalizes on SpyCas9’s uniquely stringent requirement for negative supercoils to form a productive R-loop^13–16^. Such a resistance mechanism could be accessible to diverse MGEs, providing a route to CRISPR-Cas tolerance in many genetic contexts.

## Results

### Functional selection reveals a novel anti-CRISPR protein, AcrIIA22

We recently carried out a functional selection for SpyCas9 antagonism, recovering clones from metagenomic libraries that could potently inhibit SpyCas9^12^. In this two-plasmid setup, we used an inducible SpyCas9 on an expression plasmid to cleave the *kanamycin resistance* (*Kan^R^*) gene of a second ‘target’ plasmid. We then grew cultures in SpyCas9-inducing conditions and measured the proportion of colony forming units (cfus) that remained kanamycin resistant (Figure 1A). This proportion is a measure of how many clones retained their target plasmid and thus how effectively that plasmid withstood SpyCas9 attack. In our previously published work, we describe AcrIIA11, a novel anti-CRISPR from a metagenomic clone named F01A_2 (Genbank ID MK637582.1), which was the most abundant from functional selection of a human fecal microbiome^12^. This functional selection also revealed a second protective clone, F01A_4 (Genbank ID MK637587.1). Together, these two contigs (F01A_2 and F01A_4) accounted for >96% of the normalized read coverage and were the most abundant clones recovered from this library.

**Figure 1.**
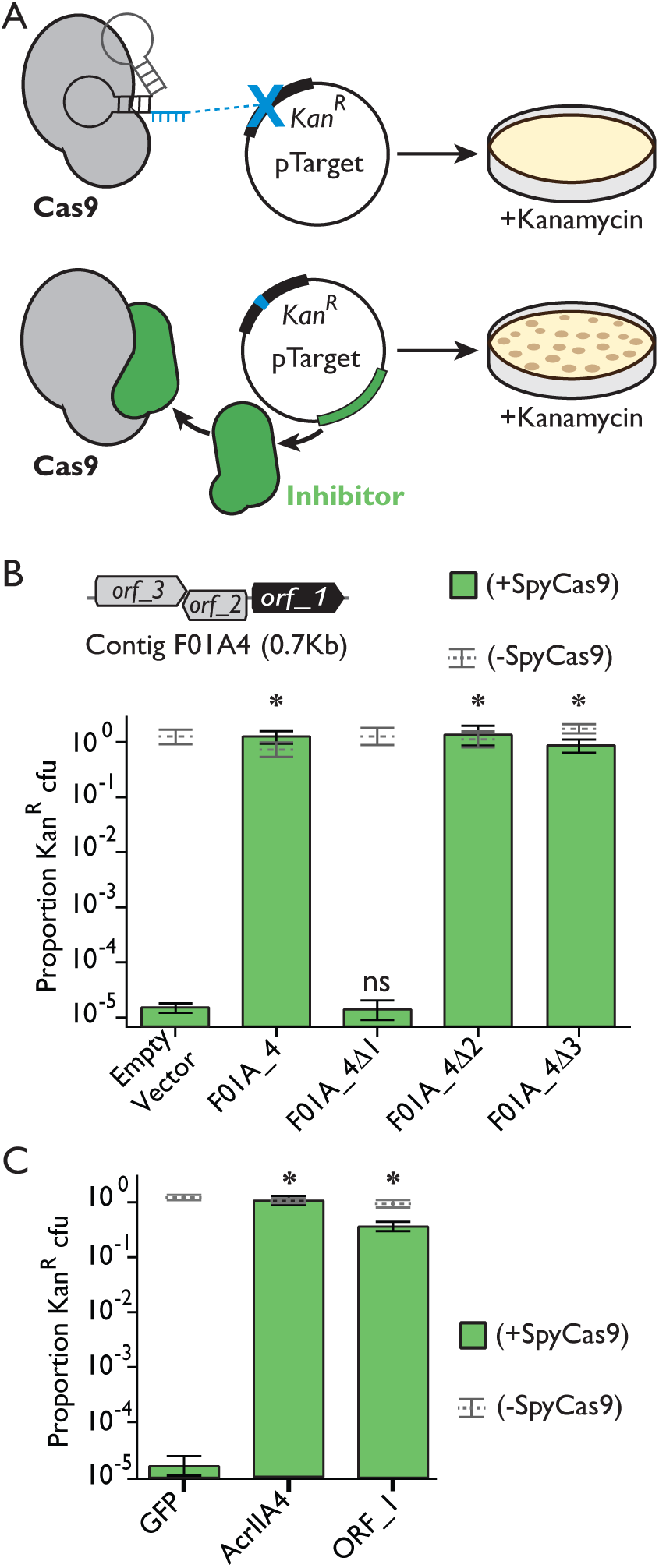
*Orf_1* from the metagenomic contig F01A_4 encodes a SpyCas9 inhibitor. (**A**) The plasmid protection assay used to reveal SpyCas9 inhibition. Plasmids without SpyCas9 inhibitors are cleaved by Cas9 and do not give rise to Kan^R^ colonies. Those with inhibitors withstand SpyCas9 attack and yield colonies. (**B**) An early stop codon in *orf_1 (Δ1),* but not *orf_2* or *orf_3 (Δ2* and *Δ3),* eliminates the ability of contig F01A_4 to protect a plasmid from SpyCas9. Asterisks depict statistically significant differences in plasmid retention between the indicated genotype and an empty vector control in SpyCas9-inducing conditions (Student’s t-test, p<0.002, n=3); ns indicates no significance. All p-values were corrected for multiple hypotheses using Bonferroni’s method. (**C**) Expression of *orf_1* is sufficient for SpyCas9 antagonism, protecting a plasmid as well as *acrIIA4*. Asterisks are as in panel B but relate to the GFP negative control rather than an empty vector.

The F01A_4 contig is 685 bp long, encodes three potential open reading frames (ORFs), and confers complete protection against SpyCas9, with plasmid retention equaling that of an uninduced SpyCas9 control (Figure 1B). To determine the genetic basis for SpyCas9 antagonism in this contig, we introduced an early stop codon into each of the three potential ORFs and analyzed how these mutations affected the contig’s ability to protect a target plasmid from SpyCas9. We found that an early stop codon in *orf_1* reduced the proportion of *Kan^R^* cfus by a factor of 10^5^, matching the value observed for an empty vector control (Figure 1B). Furthermore, expression of *orf_1* alone was also sufficient for SpyCas9 antagonism (Figure 1C), protecting a target plasmid from SpyCas9 cleavage as well as the potent SpyCas9 inhibitor, AcrIIA4. In this assay, *orf_1* was slightly toxic when singly expressed in *E. coli*, reducing growth rate by 7% (Supplemental Figure 1). Combined, our results indicate that *orf_1* completely accounts for the SpyCas9 protection phenotype of contig F01A_4.

One trivial mechanism by which *orf_1* could apparently antagonize SpyCas9 in our functional assay would be to lower its expression. To address this possibility, we carried out two experiments. First, we swapped the *spycas9* gene for *gfp* in our expression vector and asked whether *orf_1* induction impacted fluorescence output. We saw no change in fluorescence upon *orf_1* induction, indicating that *orf_1* neither suppressed transcription from our expression vector nor altered its copy number (Supplemental Figure 2). Second, we used Western blots to test whether *orf_1* expression impacted SpyCas9 protein levels through the course of a plasmid protection assay. We used a crRNA that did not target our plasmid backbone to ensure that *orf_1* expression remained high and its potential impact on SpyCas9 expression levels would be most evident. We observed that *orf_1* expression had no meaningful effect on SpyCas9 expression at any timepoint. Thus, we conclude that *orf_1* does not impact SpyCas9’s translation or degradation rate (Supplemental Figure 2). Therefore, *orf_1* must act via an alternative mechanism to inhibit SpyCas9 activity. Based on these findings, we conclude that *orf_1* encodes a *bona fide* anti-CRISPR protein and hereafter refer to it as *acrIIA22*.

Next, we investigated whether *acrIIA22* could also allow phages to escape from SpyCas9 immunity (Supplemental Figure 3). We measured SpyCas9’s ability to protect *E. coli* from infection by phage Mu, in the presence or absence of *acrIIA22*. As a control, we carried out similar phage infections in the presence or absence of the well-established SpyCas9 inhibitor, *acrIIA4*. As anticipated, SpyCas9 significantly impaired Mu when targeted to the phage’s genome but not if a non-targeting CRISPR RNA (crRNA) was used. Consistent with previous findings^12^, phage Mu could infect targeting and non-targeting strains equally well when we expressed *acrIIA4*, indicating that SpyCas9 immunity was completely abolished by this *acr*. However, *acrIIA22* could only partially restore the infectivity of phage Mu across multiple experimental conditions (Supplemental Figure 3). We therefore conclude that *acrIIA22* only weakly protects Mu phage from SpyCas9 whereas it strongly protects plasmids against SpyCas9 cleavage.

### AcrIIA22 homologs are present in hypervariable regions of bacterial and prophage genomes

AcrIIA22 is 54 amino acids in length and has no sequence homology to any protein of known function, including all previously described Acrs. We examined the distribution of *acrIIA22* homologs in NCBI’s NR and WGS databases but found just seven hits, limiting our ability to make evolutionary inferences about its origins or prevalence. We therefore expanded our search to include IMG/VR, a curated database of cultured and uncultured DNA viruses^17^, and assembly data from a meta-analysis of 9,428 diverse human microbiome samples^18^. With additional homologs from these databases in hand, we found that the majority of *acrIIA22* homologs exist in either of two genomic contexts: prophage genomes or small, bacterial genomic islands (Figures 2A, 2B). The original metagenomic DNA fragment from our selection, F01A_4, shared perfect nucleotide identity with one of these genomic islands (Figure 2B).

**Figure 2.**
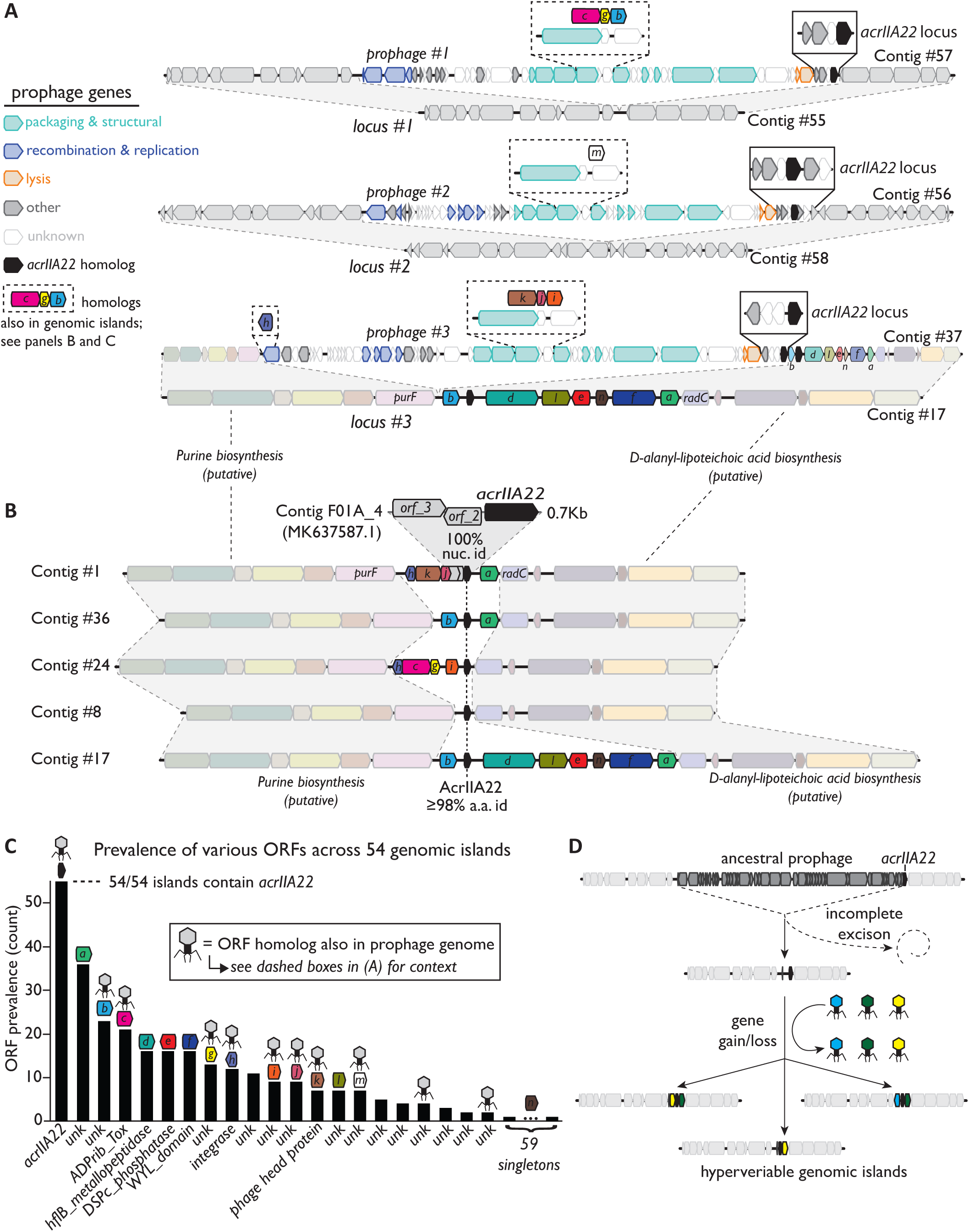
*AcrIIA22* homologs are found in hypervariable regions of prophage and bacterial genomes. (**A**) Homologs of *acrIIA22* are depicted in three related prophage genomes, integrated at three different genomic loci, revealed by a comparison of prophage-bearing contigs (#57, #56, #37) relative to unintegrated contigs (#55, #58, #17 respectively) that are otherwise nearly identical. Prophage genes are colored by functional category, according to the legend at the left of panel A. Genes immediately adjacent to *acrIIA22* (solid boxes) vary across phages, despite strong relatedness across much of the prophage genomes. Bacterial genes are colored gray, except for in contig #17, which is also depicted in panel B, below. (**B**) Homologs of *acrIIA22* are depicted in diverse genomic islands, including Contig #1, whose sequence has perfect nucleotide identity to the original metagenomic contig we recovered (F01A_4). All *acrIIA22* homologs in these loci are closely related but differ in their adjacent genes, which often have homologs in the prophages depicted in panel A (dashed boxes). Genomic regions flanking these hypervariable islands are nearly identical to one another and to prophage integration locus #3, as shown by homology to contig #17 from panel A. (**C**) The prevalence of various protein families (clustered at 65% amino acid identity) in a set of 54 unique genomic islands is shown. Each of these islands is flanked by the conserved genes *purF* and *radC* but contains a different arrangement of encoded genes. Domain-level annotations are indicated below each protein family (unk; unknown function). Gene symbols above each protein family are colored and lettered to indicate their counterparts or homologs in panels A and B. The phage capsid icon indicates sequences with homologs in prophage genomes. (**D**) An evolutionary model for the origin of the *acrIIA22*-encoding hypervariable genomic islands depicted in panel B is shown. We propose that *acrIIA22* moved via a phage insertion into a bacterial genomic locus, remained following an incomplete prophage excision event, and its neighboring genes subsequently diversified via horizontal exchange with additional phage genomes. Contigs are numbered to indicate their descriptions in Supplemental Table 3, which contains their metadata, taxonomy, and sequence retrieval information. All sequences and annotations may also be found in Supplementary Datasets 1 and 2.

Because most *acr*s are found in phage genomes, we first examined the prophages that encoded AcrIIA22 homologs. These prophages were clearly related, based on many homologous genes and a similar genome organization (Figure 2A). Despite their similarity, we found these prophages inserted into several different bacterial loci, including one site between the bacterial genes *purF* and *radC* (locus #3, Figure 2A). This prophage insertion site is notable because it is nearly identical to the highly conserved sequences that flanked *acrIIA22*-encoding bacterial genomic islands (Figure 2B). Due to their common genomic loci, we hypothesized that the apparently bacterial *acrIIA22* homologs in these genomic islands diverged from a common phage ancestor, encoded by a prophage that previously integrated at this locus. We speculate that the original *acrIIA22*-encoding bacterial genomic island was left behind following the incomplete excision of an ancestral, *acrIIA22*-encoding prophage. Supporting this hypothesis, *acrIIA22* homologs are always found at the end of prophage genomes, near their junction with a host bacterial genome (Figure 2A).

To better understand *acrIIA22*’s gene neighborhood, we again searched the raw assemblies of over 9,400 human microbiomes for more examples of these genomic islands^18^, but did not include *acrIIA22* in our second search criteria. Instead, we focused on the recent evolutionary history of these bacterial genomic islands by only considering contigs with ≥98% nucleotide identity to *purF* and *radC*, the conserved genes that flanked the genomic islands. This search yielded 258 contigs. Aligning these sequences revealed that each contig encoded a short, hypervariable region of small ORFs which was flanked by conserved genomic sequences (Figure 2B). In total, we observed 128 unique examples of these hypervariable loci, which displayed considerable gene turnover, resulting in 54 distinct gene arrangements among the 128 unique loci. Despite not including them in our search strategy, *acrIIA22* homologs were universally conserved in all 128 unique genomic islands whereas no other gene was present in more than two-thirds of the 54 distinct gene arrangements (Figure 2C). Based on this finding, we infer that the arrival of *acrIIA22* preceded the diversification seen at this locus and that its homologs have been retained since, despite the considerable gene turnover that has occurred subsequently.

Though most ORFs in these islands were of unknown function, many had close homologs in the genomes of nine representative *acrIIA22*-encoding phage (dashed boxes in Figure 2A, phage icons in Figure 2C). This suggests that phages continue to supply the genetic diversity seen at these hypervariable genomic loci. These rapid gene gains and losses probably occur as they do in other genomic islands, via recombination between this locus and related MGEs that infect the same host bacterium without the MGE necessarily integrating into the locus^19^. Taken together, our data suggest that an incomplete prophage excision event left *acrIIA22* behind in a bacterial genomic locus, which then diversified via gene exchange with additional phage genomes (Figure 2D).

In prophage genomes, *acrIIA22* homologs were found in hypervariable regions, near the junction with the host bacterial genome (Figure 2A). Both features imply these loci are subject to higher than average rates of recombination. Despite this, we could find no gene consistently present within or outside of these genomic islands that could account for their hypervariable nature (*e.g.* an integrase, transposase, recombinase, or similar function that is typically associated with genomic islands^20^). Instead, *acrIIA22* was the only gene conserved at this locus. If it could somehow promote recombination, either alone or with other factors, this could account for the high rates of gene exchange observed adjacent to *acrIIA22* in phage and bacterial genomes (Figures 2A, 2B).

In total, we identified 30 unique *acrIIA22* homologs, 25 of which were predicted to originate from the unnamed *Clostridial* genus, CAG-217 (Figure 3A). Because *acr*s are only beneficial to phages if they inhibit CRISPR-Cas activity, they are typically found only in taxa with a high prevalence of susceptible Cas proteins^9^. If AcrIIA22 functions naturally as an Acr, we would predict that Cas9-encoding, type II-A CRISPR-Cas systems like SpyCas9 would be common in CAG-217 bacteria. To test this idea, we examined 779 draft assemblies of CAG-217 genomes and found that 179 of the 181 predicted CRISPR-Cas systems in CAG-217 genomes were Cas9-encoding, type II-A systems. This enrichment for Cas9 is particularly striking for a *Clostridial* genus, as *Clostridia* rarely encode Cas9. Instead, they typically encode other CRISPR-Cas defenses^21^. Thus, the distribution of CRISPR-Cas systems in CAG-217 genomes supports our hypothesis that *acrIIA22* functions natively as an *acr*.

**Figure 3.**
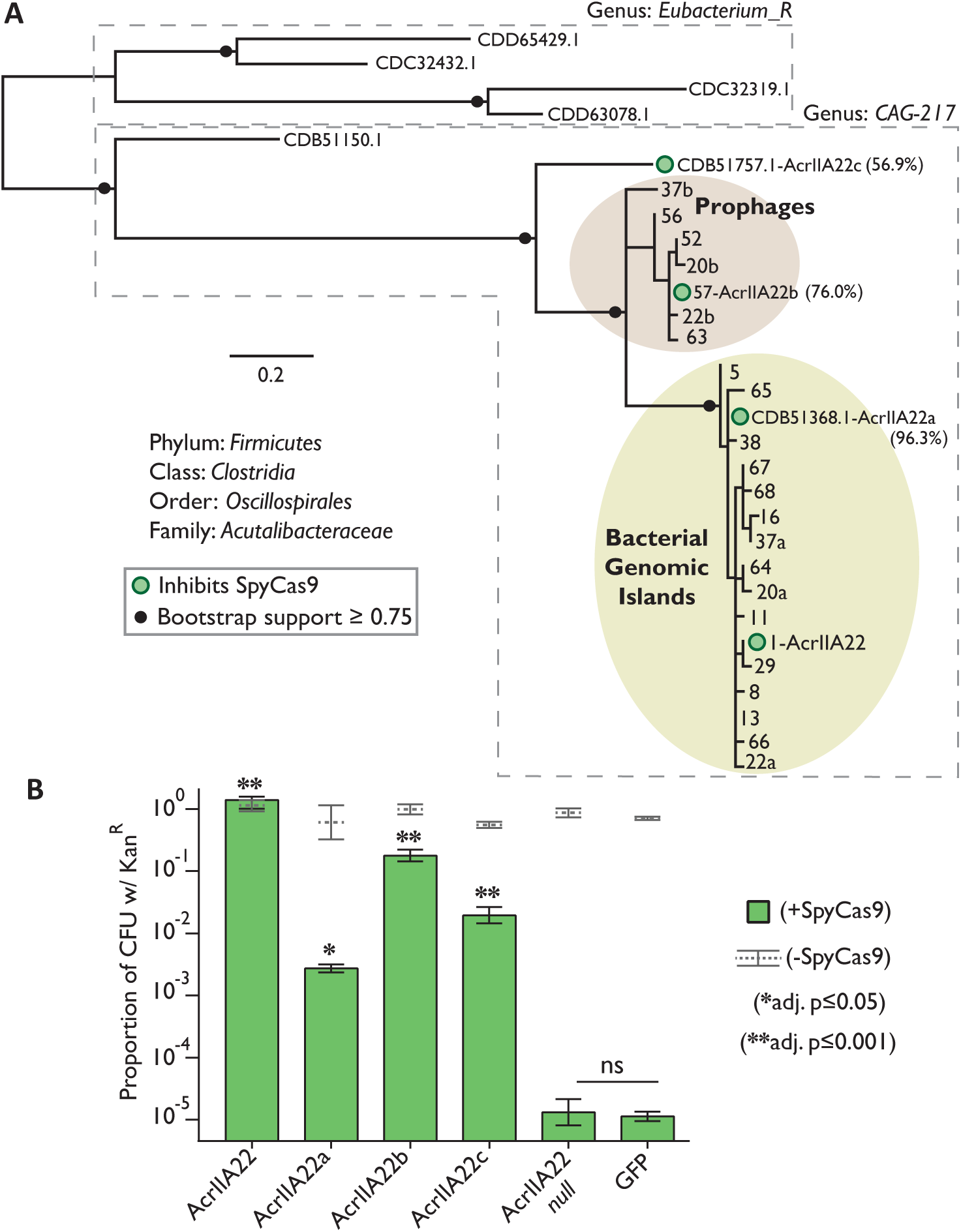
AcrIIA22 homologs capable of inhibiting SpyCas9 are common in the unnamed *Clostridial* genus, CAG-217. Phylogenetic classifications were assigned corresponding to the GTDB naming convention (Methods). (**A**) A phylogeny of all unique AcrIIA22 homologs identified from metagenomic and NCBI databases. Prophage sequences are shaded brown and homologs from hypervariable bacterial genomic islands are shaded yellow. Sequences obtained from NCBI are labeled with protein accession numbers. In other cases, AcrIIA22 homologs are numbered to match their contig-of-origin (Supplemental Table 3). In some cases, more than one AcrIIA22 homolog is found on the same contig (‘a’ or ‘b’ indicates its presence in a hypervariable genomic island or prophage genome, respectively). Circles at nodes indicate bootstrap support ≥ 0.75. Dashed boxes separate sequences identified from different bacterial genera. Filled green circles indicate homologs that were tested for their ability to inhibit SpyCas9 in the plasmid protection assay in panel B. (**B**) Homologs of AcrIIA22 in CAG-217 genomes inhibit SpyCas9. Asterisks depict statistically significant differences in plasmid retention under SpyCas9-inducing conditions between the indicated sample and a null mutant with an early stop codon in *acrIIA22*, per the legend at right (ns indicates no significance). All p-values were corrected for multiple hypotheses using Bonferroni’s method (Student’s t-test, n=3).

We also found evidence that Cas9 is active in CAG-217 bacteria. Prophages from CAG-217 encode 78 type II-A Acrs (homologs of AcrIIA7, AcrIIA17, and AcrIIA21), suggesting they are actively engaged in an arms race with Cas9-based defenses in these bacteria. We even found one example where homologs of *acrIIA17* and *acrIIA22* were located within one kilobase of each other in a prophage genome (Supplemental Figure 4)^22^. Since phages often aggregate *acr*s in the same genomic locus^23^, this observation independently supports our hypothesis that CAG-217 prophages encode *acrIIA22* homologs to inhibit type II-A CRISPR-Cas systems.

We next tested whether the ability to inhibit type II-A CRISPR-Cas systems was a shared property of *acrIIA22* homologs from CAG-217 bacteria. To do so, we selected *acrIIA22* homologs that spanned the phylogenetic diversity present among CAG-217 genomes (Figure 3A) and tested their ability to protect a target plasmid from SpyCas9 elimination. These analyses revealed that each *acrIIA22* homolog from CAG-217 could antagonize SpyCas9 activity at least partially (Figure 3B). This conservation of anti-SpyCas9 activity among divergent AcrIIA22 homologs (for example, sharing only 56.9% identity), suggests that they may broadly inhibit Cas9. Broad inhibition has been seen for some other type II-A Acrs^12^ and can occur either by targeting a conserved feature of Cas9 or by inhibiting Cas9 via an indirect mechanism that it cannot easily evade.

### AcrIIA22 functions via a non-canonical mechanism

Almost all characterized Acrs inhibit their cognate Cas proteins via direct binding without the involvement of additional co-factors; as a result, they exhibit strong inhibitory activity when tested *in vitro* (Supplemental Table 1). To determine if this was the case for AcrIIA22, we purified it from *E. coli* and asked whether it could bind and inhibit SpyCas9. To test for binding, we asked whether a tagged AcrIIA22 co-precipitated with SpyCas9 when mixed as purified proteins. Unlike AcrIIA4, which binds strongly to SpyCas9 and inhibits its activity *in vitro*, we could detect little to no binding between AcrIIA22 and SpyCas9, regardless of whether a single-guide RNA (sgRNA) was included or not (Supplemental Figure 5). We also observed that AcrIIA22 had no impact on SpyCas9’s ability to cleave linear, double-stranded DNA (dsDNA), even when AcrIIA22 was included at substantial molar excess over SpyCas9 (Supplemental Figure 6). These results suggest that, at least in isolation, AcrIIA22 cannot bind and inhibit SpyCas9. Thus, AcrIIA22 lacks the predominant biochemical activities exhibited by previous Acrs that have been mechanistically characterized.

We therefore considered the possibility that AcrIIA22 encodes an unconventional anti-CRISPR that acts via a non-canonical mechanism. However, AcrIIA22 homologs had no sequence homology to other characterized proteins, which would have provided clues about AcrIIA22 activity and biochemical mechanisms. Anticipating that structural homology might provide some insight, we solved AcrIIA22’s structure using X-ray crystallography. We first built a homology model from AcrIIA22’s primary sequence with Robetta. We then used this model for molecular replacement to solve its structure at 2.80Å resolution (PDB:7JTA). The asymmetric unit in AcrIIA22’s crystal comprises two monomers stacked end-to-end, with each monomer folding into a four-stranded β-sheet (Figure 4A, Table 1). A DALI structure-structure search revealed that the AcrIIA22 monomer is similar to members of the newly recognized PC4-like structural fold (Figure 4B, Supplemental Table 2). PC4-like proteins have independently evolved in all domains of life, typically adopt a β-β-β-β-α topology, and often homodimerize to bind diverse RNA and DNA species using variably positioned β-sheets^24^.

**Figure 4.**
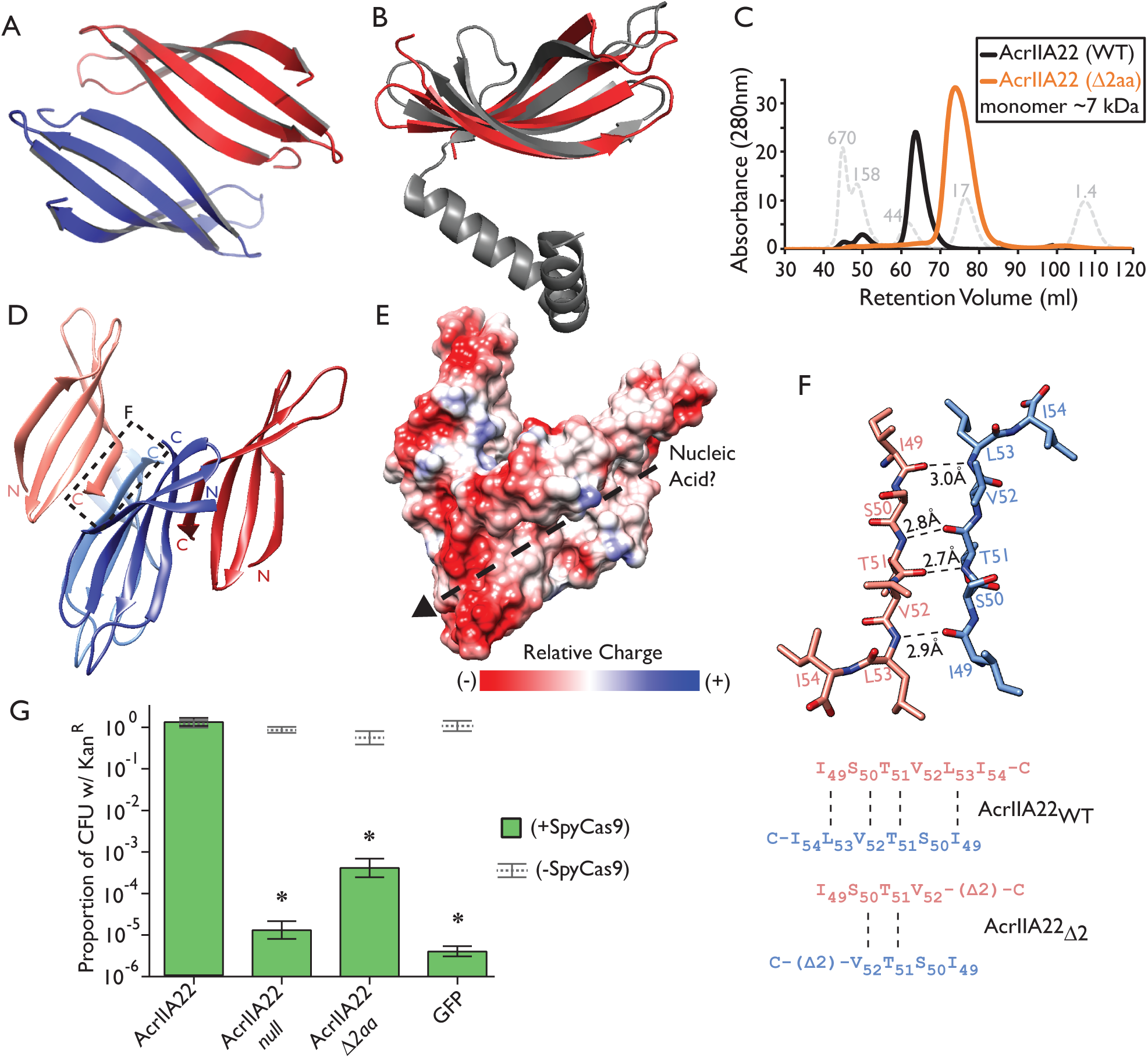
AcrIIA22 is a PC4-like protein that oligomerizes and inhibits SpyCas9. (**A**) AcrIIA22’s crystal structure reveals a homodimer of two four-stranded β-sheets. (**B**) A monomer of AcrIIA22 (PDB:7JTA) is structurally similar to a predicted single-stranded DNA binding protein, which is proposed to promote recombination in phage T5 (PDB:4BG7, Z-score=6.2, matched residues 15%). (**C**) AcrIIA22 elutes as an oligomer that is 4-5 times the predicted molecular mass of its monomer. The gray, dashed trace depicts protein standards of the indicated molecular weight. The orange trace depicts the elution profile of a two-amino acid C-terminal AcrIIA22 truncation mutant. (**D**) Ribbon diagram of a proposed AcrIIA22 tetramer which requires binding between anti-parallel β-strands at the C-termini of AcrIIA22 monomers to form extended, concave β-sheets. This putative oligomerization interface is indicated by the dashed box and is detailed in panel F. (**E**) Space filling model of the tetrameric AcrIIA22 structure from panel D, with relative charge depicted, highlighting a groove (dashed line with arrowhead) that may accommodate nucleic acids. (**F**) A putative oligomerization interface between the C-termini of two AcrIIA22 monomers is shown with hydrogen bond distances between the polypeptide backbones indicated. The wild-type sequence and truncation mutant are indicated below. Dashed lines indicate potential hydrogen bonds. This interface occurs twice in the putative tetramer, between red-hued and blue-hued monomers in panel D. (**G**) The truncation mutant fails to protect a plasmid from SpyCas9 elimination, similar to an early stop codon mutant (*null*) and a *gfp* negative control. Asterisks depict statistically significant differences in plasmid retention under SpyCas9-inducing conditions between the indicated sample and the wild-type sequence (adj. p < 0.002, Student’s t-test, n=3). All p-values were corrected for multiple hypotheses using Bonferroni’s method.

**Table 1.** Structural features of AcrIIA22.

| **Data collection** |  |
| --- | --- |
| Space Group | P4332 |
| *Cell Dimensions* |  |
| a, b, c (Å) | 128.56, 128.56, 128.56 |
| $\alpha$, $\beta$, $\gamma$ (°) | 90.0, 90.0, 90.0 |
| Resolution (Å) | 50.00 - 2.80 |
| $R_{merge}$ | 0.106 (0.906) |
| $I/\sigma_I$ | 17.4 (2.6) |
| Completeness (%) | 98.7 (100.0) |
| Redundancy | 10.4 (10.7) |
| CC 1/2 | 0.837 |
| **Refinement** |  |
| No. Reflections | 9334 |
| $R_{work}$ ($R_{free}$) (%) | 22.2 (24.6) |
| No. Complex in ASU | 2 |
| *No. atoms* |  |
| Protein | 810 |
| Heteroatoms | 50 |
| Water | 3 |
| B-factor | 82.82 |
| *R.m.s deviations* |  |
| Bond lengths (Å) | 0.003 |
| Bond angles (°) | 0.610 |
| *Ramachandran* |  |
| Preferred (%) | 98.15 |
| Allowed (%) | 1.85 |
| Outliers (%) | 0 |

Despite crystallizing as a homodimer, AcrIIA22 migrated at a size substantially larger than its molecular weight by size exclusion chromatography (SEC) (Figure 4C). This suggested that AcrIIA22 may oligomerize *in vivo*. Structural evidence supported the same conclusion, as AcrIIA22 was predicted to form a stable tetramer when analyzed with PISA, a tool for inferring macromolecular assembles from crystalline structure^25^ (Figures 4D, 4E). This putative tetramer has a molecular mass consistent with that observed by SEC and comprises pairs of outward-facing, concave β-sheets, as is seen in other PC4-like proteins^24^. Interestingly, many PC4-like proteins bind nucleic acids using similar concave β-sheets and, in some instances, form higher-order oligomers as a necessary step for binding DNA or RNA^24^. Consistent with this possibility, adjacent β-sheets along each outward face of the putative AcrIIA22 tetramer form a groove that could potentially accommodate a nucleic acid substrate (Figure 4E). Thus, even though AcrIIA22 lacks the alpha-helix typically seen in PC4-like proteins, its other structural and functional attributes led us to suspect that AcrIIA22 also interacted with nucleic acids.

Our tetramer model predicts that a four amino acid interface at the C-terminus of AcrIIA22 is required for adjacent β-sheets to bind one another and form a grooved, oligomeric structure (Figures 4D, 4F). We predicted that a two-residue, C-terminal truncation of AcrIIA22 would disrupt this interface (Figure 4F). To test this prediction, we examined the oligomeric state of this 2-aa AcrIIA22 deletion mutant. Consistent with our hypothesis, we found that these AcrIIA22 complexes migrated at about half the size of their wild-type counterparts by SEC (Figure 4C), suggesting that this C-terminal interface is required to progress from a two to four-membered oligomer. Moreover, we found that the 2-aa deletion mutant was also impaired for SpyCas9 antagonism in our plasmid protection assay (Figure 4G). Thus, this C-terminal motif is necessary for protection from SpyCas9 and higher-order oligomerization, suggesting that oligomerization may be necessary for AcrIIA22’s anti-SpyCas9 activity.

### AcrIIA22 is a DNA nickase that relieves torsion of supercoiled plasmids

Our structural analyses indicated that AcrIIA22 is a PC4-like nucleic acid-interacting protein. Like AcrIIA22, many PC4-like proteins are encoded in phage genomes. Among these is AcrIIA22’s closest structural relative in the PC4 family: a predicted single-stranded binding (SSB) protein from phage T5 (Figure 4B)^26^. This putative SSB protein has been predicted to directly stimulate recombination during the recombination-dependent replication of phage T5’s genome^27^. This prediction, together with our inference from genomic analyses (Figure 2), led us to hypothesize that AcrIIA22 may have similar recombination-stimulating activity. Indeed, other PC4-like proteins have been observed experimentally to unwind duplex DNA, a function consistent with their proposed roles in transcription and recombination^24,28^. Therefore, we investigated whether AcrIIA22 might interact with duplexed DNA in a manner consistent with its putative recombinogenic properties.

We first asked whether we could detect any biochemical effect of *acrIIA22* on a double-stranded DNA (dsDNA) target plasmid *in vivo*. In this experiment, we considered three *acrIIA22* genotypes: the wild-type sequence, a null mutant with a single base pair change to create an early stop codon, and the 2-aa truncation mutant that we previously showed was defective for oligomerization (Figure 4C) and SpyCas9 antagonism (Figure 4G). We then grew overnight cultures of plasmids expressing each genotype, purified plasmid DNA, and analyzed its topology using gel electrophoresis (Figure 5A). As is typical for plasmid purifications from *E. coli*, the plasmid encoding the null mutant was predominantly recovered in a supercoiled form. In contrast, AcrIIA22 expression shifted much of the target plasmid to a slowly migrating form, consistent with an open-circle conformation. These findings suggest that AcrIIA22 expression could relieve plasmid supercoiling, hinting at a potential DNA nickase activity. We also found that the 2-aa truncation mutant was impaired for this putative nickase activity, consistent with this mutant’s compromised oligomerization and anti-Cas9 activities (Figure 4G).

**Figure 5.**
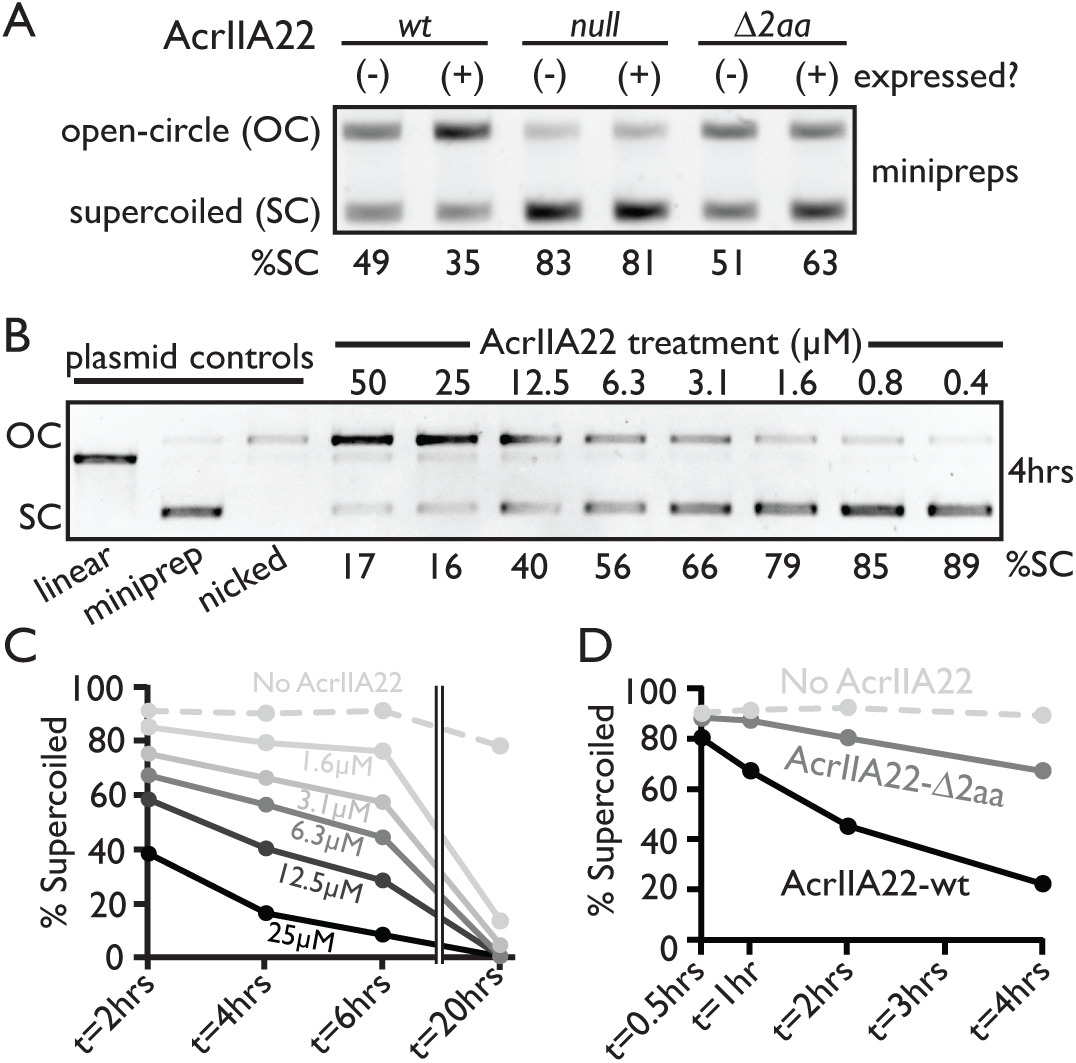
AcrIIA22 nicks supercoiled plasmids *in vivo* and *in vitro*. (**A**) Gel electrophoresis of plasmids purified from overnight *E. coli* cultures expressing the indicated genotypes. OC, open-circle plasmid; SC, supercoiled plasmid. %SC indicates the percentage of DNA in the supercoiled form for each sample. (**B**) AcrIIA22 nicks supercoiled plasmids *in vitro*. Supplemental Figure 7 depicts this experiment at additional time points. (**C**) Quantification of AcrIIA22-nicked plasmids in panel B and Supplemental Figure 7. AcrIIA22 nicks plasmids in a time and concentration-dependent manner. (**D**) A nickase assay as in panels B and C shows that the 2-aa truncation mutant is impaired for activity *in vitro*, relative to wild-type AcrIIA22. In both cases, 25 µM protein was used.

Because *acrIIA22* expression altered plasmid topology *in vivo*, we next asked whether purified AcrIIA22 had an impact on a plasmid DNA substrate *in vitro*. By gel electrophoresis, we observed that AcrIIA22 shifted a supercoiled plasmid to a slowly migrating form in a time and concentration-dependent manner (Figures 5B, 5C, Supplemental Figure 7). For comparison, we also treated a plasmid with the nickase Nb.BssSI, yielding a band that migrated at the same position as the putatively open-circle product generated via AcrIIA22 activity (Figure 5B). Extended incubation times and high concentrations of AcrIIA22 resulted in conversion of plasmids to a linearized DNA product, consistent with a nickase-like nuclease activity acting on both strands of DNA (Figure 5B, Supplemental Figure 7). AcrIIA22’s nickase activity was strongly stimulated in the presence of Mn^2+^, Co^2+^, and Mg^2+^, weakly with Ni^2+^ and Zn^2+^, but not at all with Ca^2+^ (Supplemental Figure 8). Consistent with our *in vivo* observations, we found that the 2-aa deletion mutant was impaired for nickase activity, relative to wildtype AcrIIA22 (Figure 5D).

Our *in vitro* and *in vivo* findings suggest that AcrIIA22 is responsible for the change in plasmid topology observed in bacterial cells. To confirm that the observed gel-shift was the result of AcrIIA22 nickase activity and not protein-bound DNA, we purified an AcrIIA22-treated plasmid with phenol-chloroform and re-examined it by gel electrophoresis. We observed that the nicked form of the plasmid persisted through purification, establishing AcrIIA22 as a *bona-fide* nickase (supp figure 8). Therefore, based on both *in vitro* and *in vivo* findings, we conclude that that *acrIIA22* encodes a nickase protein that relieves the torsional stress of supercoiled plasmids.

### AcrIIA22’s nickase activity impairs SpyCas9

Having established that AcrIIA22 is a DNA nickase, we next investigated whether this biochemical activity correlated with its ability to inhibit SpyCas9. If this were the case, it would explain how AcrIIA22 protected plasmids from SpyCas9 without directly binding the Cas protein. We therefore tested the consequences of expressing AcrIIA22 on a target plasmid in the presence of SpyCas9. As before, we began by comparing overnight plasmid purifications of a target plasmid expressing AcrIIA22, a null mutant with an early stop codon, or the 2-aa AcrIIA22 truncation mutant. However, this time, we also subjected the plasmid to SpyCas9 targeting during bacterial growth. We were unable to recover the negative control target plasmid after overnight growth, implying that this target plasmid was eliminated by SpyCas9 (Figure 6A). The 2-aa truncation mutant was also eliminated by SpyCas9, indicating these residues are important for function. In contrast, SpyCas9 did not eliminate a target plasmid that expressed full-length AcrIIA22 (Figure 6A), consistent with AcrIIA22’s previously established capacity to protect against SpyCas9 (Figure 1C).

**Figure 6.**
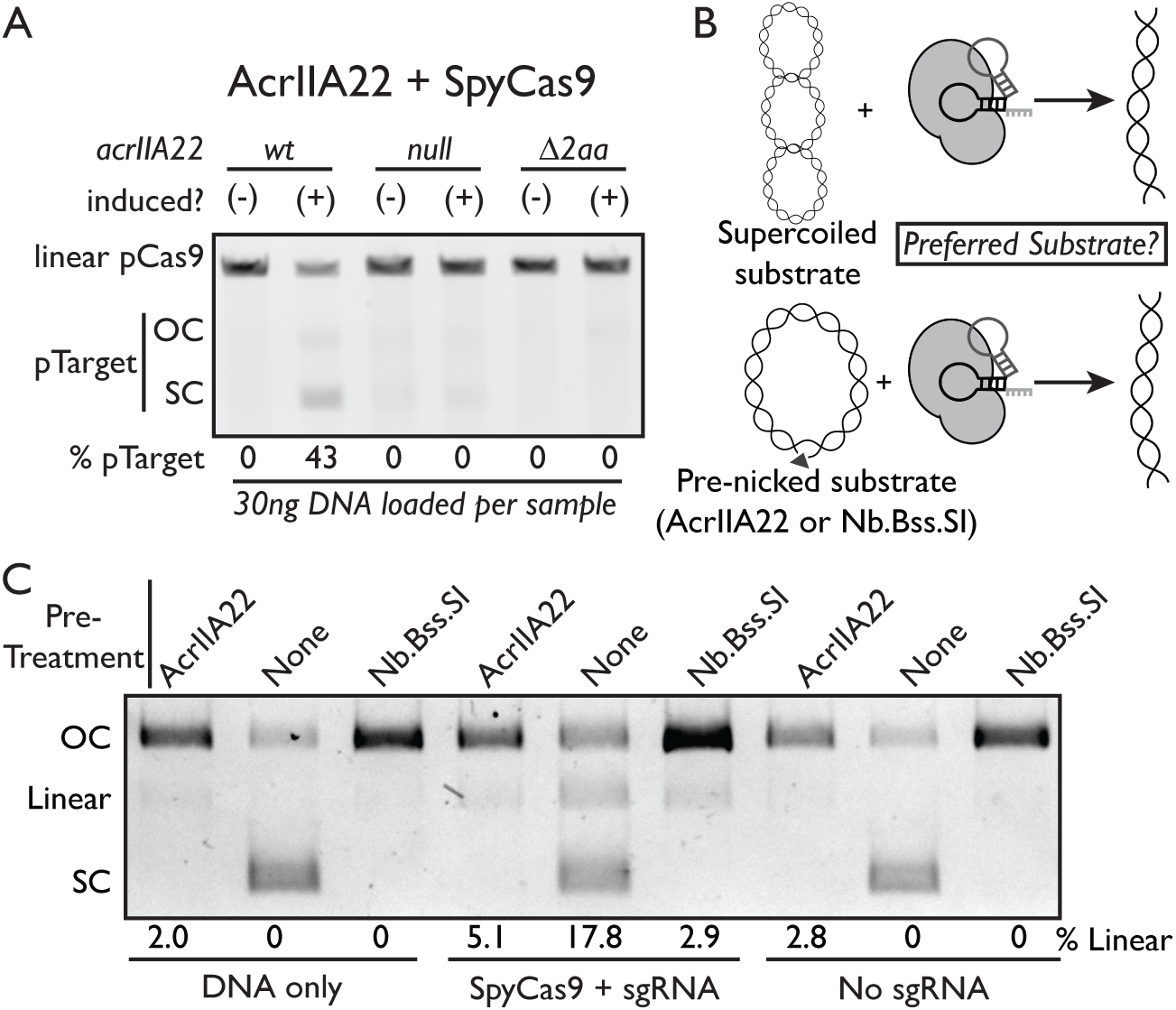
AcrIIA22 protects plasmids from SpyCas9 cleavage *in vivo* and *in vitro*. (**A**) Gel electrophoresis of plasmids purified from overnight *E. coli* cultures expressing the indicated *acrIIA22* genotypes and SpyCas9 from a second plasmid. The *acrIIA22*-encoding plasmids are indicated with the ‘pTarget’ label. OC, open-circle; SC, supercoiled. The SpyCas9 plasmid was linearized via a unique restriction site before electrophoresis. (**B**) Experimental design for the data depicted in panel C. The experiment tests whether SpyCas9 preferentially cleaves a supercoiled or open-circle plasmid target *in vitro*. (**C**) Nicked plasmids are less susceptible to linearization via SpyCas9 cleavage. Plasmid purifications from overnight cultures were either left unmodified or pre-treated with one of two nickase enzymes, AcrIIA22 or Nb.Bss.SI. Linear, open-circle (OC), and supercoiled (SC) plasmid forms are indicated. The percentage of DNA in the linear form is quantified below the gel, where the reaction components are also listed. See Supplemental Figure 9 for these data in different reaction conditions.

To be effective, a CRISPR-Cas system must eliminate its target at a faster rate than the target can replicate^29^. Our findings raised the possibility that AcrIIA22 modifies a target plasmid into a SpyCas9-resistant conformation to win this ‘kinetic race’ against SpyCas9, potentially shifting the equilibrium to favor plasmid persistence instead of elimination. To test this kinetic race model, we asked whether a plasmid that had been pre-treated with AcrIIA22 could resist digestion by SpyCas9 *in vitro*. Therefore, we purified the open-circle plasmid that resulted from AcrIIA22 pre-treatment and determined how efficiently it was cleaved by SpyCas9 compared to an unmodified, supercoiled plasmid (Figure 6B). SpyCas9 showed a clear preference for cleaving the supercoiled substrate versus the AcrIIA22-treated open-circle plasmid (Figure 6C, Supplemental Figure 9). An open-circle plasmid pre-treated with the nickase Nb.Bss.SI was similarly recalcitrant to SpyCas9 digestion. Taken together, our findings suggest that relieving DNA torsion provides the mechanistic explanation for AcrIIA22’s ability to inhibit SpyCas9 *in vivo*.

Our findings also help explain why AcrIIA22 is more adept at protecting plasmids than phages from SpyCas9 in our system. Because plasmids are maintained as circular, extrachromosomal elements, they are more likely to undergo torsional change when nicked than phages or transposons, which are often linear or spend significant time integrated into their host’s genome. Additionally, linear DNA experiences minimal torsional stress and is therefore less susceptible than supercoiled plasmids to cleavage by SpyCas9^15^. This likely explains why AcrIIA22 failed to protect a linear dsDNA substrate from SpyCas9 *in vitro* (Supplemental Figure 6), as there is very little torsional stress for it to relieve in this substrate. Importantly, *in vitro* experiments indicate that Cas9 requires a higher degree of negative supercoiling to provide the free energy needed for R-loop formation than type I CRISPR-Cas systems^13^. *In vivo* observations also show that DNA supercoiling affects the recruitment of SpyCas9 to its target site in bacteria^14^. This suggests that Cas9 may be particularly susceptible to changes in DNA torsion. Thus, factors like the AcrIIA22 nickase, which modify DNA torsion, may provide a general means to protect against Cas9.

## Discussion

In this study, we identify and characterize *acrIIA22*, a previously undescribed gene that can antagonize SpyCas9. We show that AcrIIA22 homologs have proliferated in genomes of CAG-217 bacteria, which have a high prevalence of Cas9 homologs. Using a combination of structural and biochemical studies, we show that AcrIIA22 acts by nicking supercoiled DNA to relieve torsional stress on a target plasmid, thereby impairing SpyCas9 activity *in vivo* and *in vitro*. Taken together, our data suggest that DNA topology represents a new battleground in the evolutionary arms race between CRISPR-Cas systems and MGEs. Because Cas9 is more susceptible to evasion via changes to DNA topology than other CRISPR-Cas systems^13^, it may be more disadvantaged than other bacterial defense systems in this arms race. Additionally, DNA topology is dynamically regulated in phages, plasmids and other MGEs. This means that topology-modifying factors already exist in diverse MGEs that could have secondary effects on CRISPR-Cas activity and thus prove useful in the context of a molecular arms race^30,31^. For instance, though not studied in the context of bacterial defense systems, the fitness of phage T4 is improved via the expression of an accessory protein that modifies DNA supercoiling and the propensity of R-loops to form^32^. Other phages, such as those in the T5-like family, incorporate regular nicks into their genome, the function of which has eluded description for over 40 years^33^. Based on our findings, we hypothesize that phages and MGEs targeted by Cas9 exploit factors that modify DNA topology as a tactic to evade host immunity.

Functional selections like ours are biased towards identifying genes that work well in a heterologous context. For example, even though AcrIIA22 is encoded on the genome of a genetically intractable bacterium, we could identify it using a functional metagenomic selection for SpyCas9 antagonism in *E. coli*. Although we have characterized its activities in *E. coli* and *in vitro*, it remains formally possible that AcrIIA22 functions differently in its native context. For instance, we cannot rule out the possibility that AcrIIA22 might interact with a Cas9 protein from a CAG-217 bacterium. Alternatively, AcrIIA22’s anti-Cas9 activity might be related to a native recombinogenic function (Figure 2). As precedent for this idea, CRISPR-Cas evasion was recently demonstrated for homologs of the recombination proteins Redβ and λExo in *Vibrio cholerae*^34^. Nevertheless, the heterologous behavior of AcrIIA22 in *E. coli* is clearly sufficient for SpyCas9 antagonism *in vivo* and its nickase activity can protect plasmids from SpyCas9 *in vitro*. Furthermore, an AcrIIA22 mutant that is partially defective for nickase activity *in vitro* (Figure 5D) is ∼1,000-fold less effective at protecting a plasmid from SpyCas9 *in vivo* (Figure 4G). This indicates that modest changes in nickase activity can have major consequences for plasmid survival, which is consistent with our kinetic race model (Figure 6B) and previous observations that non-linear equilibrium dynamics determine whether an MGE withstands CRISPR-Cas immunity^29^.

Our results suggest that proteins that affect DNA torsion may also enable Cas9 antagonism. For example, in addition to AcrIIA22, the Nb.BssSI nickase was also capable of protecting a plasmid from SpyCas9 *in vitro*. Yet, despite the regular occurrence of nickases in nature, functional selections for anti-Cas9 activity have not previously recovered these enzymes^12,35^. We speculate that AcrIIA22 treads a fine balance between activity and toxicity; its nickase activity is high enough to antagonize SpyCas9 in a kinetic race, but not so high that it would be toxic to the host cell (Supplemental Figure 1). Its oligomerization may represent an important mechanism to control nickase activity and suppress host toxicity. Studies of other phage- and bacterial-encoded nickase proteins may provide additional insight into whether AcrIIA22 proteins have additional properties that render them to be especially well-suited to antagonize SpyCas9.

Is AcrIIA22 a true anti-CRISPR? AcrIIA22 lacks features that are typical of conventional Acrs, such as the ability to bind Cas proteins or to inhibit CRISPR-Cas activity as a purified protein. However, other Acr proteins also lack these features. For example, the well-characterized SpyCas9 antagonist AcrIIA1 does not inhibit purified SpyCas9, but instead stimulates Cas9 degradation^36^. Similarly, AcrIIA7 does not appear to bind SpyCas9 but can nevertheless inhibit it *in vitro* via an unknown mechanism^35^. Indeed, anti-CRISPR proteins are defined by a common strategy and outcome rather than by a common biochemical mechanism. Our finding that *AcrIIA22* is encoded by prophages as a single gene that strongly protects plasmids and weakly protects phages from SpyCas9 (Figure 3B, Supplemental Figure 4) makes it much more similar to other Acrs^23^ and distinct from non-canonical CRISPR-Cas evasion strategies like DNA glucosylation^6^ or homologous recombination^34^.

AcrIIA22 does not appear to provide the same potency of Cas9 inhibition as some other characterized Acrs, particularly in protecting phage Mu. However, potent inhibition is not a pre-requisite for effective anti-CRISPR activity. Indeed, selection can favor weak anti-CRISPRs over strong ones in mixed phage populations^37^. Even in cases where mechanisms for Cas9 evasion are weak (Supplemental Figure 3), they may nonetheless confer substantial benefit. For example, slowing down Cas9 cleavage could increase the time and probability for escape mutants to arise (*e.g.* Cas9 target-site variants^1^, deletion mutants^34^), allow for additional Acr expression^38,39^, or permit further genome replication to overwhelm CRISPR-Cas immunity^29^. This phenomenon – weak tolerance giving rise to long-term resistance – is reproducibly observed in cases of strong selective pressure. For instance, in the context of antibiotic resistance, the expression of QNR pentapeptide proteins in many human pathogens can provide low-level drug tolerance, extend survival, and allow time for additional mutations to develop that completely resist quinolone antibiotics^40^.

As the use of functional metagenomics to study phage-bacterial conflicts grows more common, many novel genes and mechanisms for CRISPR-Cas inhibition are likely to be described^12,35^. Like AcrIIA22, which has no homology to any previously described anti-CRISPR and lacks other genetic signatures used for *acr* discovery (*e.g.,* linkage with helix-turn-helix transcription factors)^41,42^, these new genes may not exhibit canonical Acr behaviors. It is inevitable that these discoveries will lead a more nuanced understanding of the arms race between CRISPR-Cas systems and MGEs. While they might blur the precise boundaries on what defines an anti-CRISPR, these findings will also reveal undiscovered strategies for molecular antagonism and new battlegrounds in the age-old conflict between bacteria and their phages.

## Methods

### Plasmid protection assay

All assays were done in *Escherichia coli* (strain: NEB Turbo). As described previously^12^, SpyCas9 was expressed from a CloDF13-based plasmid marked with a spectinomycin resistance cassette. The SpyCas9 construct programed to eliminate a kanamycin-marked target plasmid was called pSpyCas9_crA (Supplemental Table 4). It eliminated a target vector that inducibly expressed a gene-of-interest via depression of the TetR transcription factor with doxycycline (named generically pZE21_tetR; Supplemental Table 4). IPTG was used in samples with the target vector to ensure high levels of TetR expression (which was driven by the lac promoter) and thus inducible control of our gene of interest. Cultures of each sample were grown overnight at 37C with shaking at 220 rpm in lysogeny broth (LB; 10 g/L casein peptone, 10 g/L NaCl, 5 g/L ultra-filtered yeast powder) containing spectinomycin 50 µg/ml, kanamycin 50 µg/ml, and 0.5mM IPTG. These growth conditions kept both SpyCas9 and the gene of interest in uninduced states. The next morning, overnight cultures were diluted 1:50 into LB broth containing spectinomycin 50 µg/ml, kanamycin 50 µg/ml, 0.5mM IPTG, and doxycycline 100 ng/ml to induce the gene of interest. Cultures were grown at 37C on a roller drum to mid-log phase (for approximately 1.5 hours to OD600 of 0.3-0.6). Once cells reached mid-log phase, they were diluted to OD600 value of 0.01 into two media types: (a) LB containing spectinomycin 50 µg/ml, 0.5mM IPTG, and doxycycline 100 ng/ml, and (b) LB containing spectinomycin 50 µg/ml, 0.5mM IPTG, doxycycline 100 ng/ml, and 0.2% (L) arabinose. These media induced either the gene of interest alone, or both the gene of interest and SpyCas9, respectively. Each sample was grown in triplicate in a 96 well plate in a BioTek Cytation 3 plate reader. After 6 hours of growth at 37°C with shaking at 220 rpm, each sample was diluted ten-fold and plated on two types of media: (a) LB spectinomycin 50 µg/ml + 0.5mM IPTG or (b) LB spectinomycin 50 µg/ml, kanamycin 50 µg/ml, 0.5mM IPTG. Plates were incubated at 37C overnight. Then, colonies were counted to determine the fraction of colony forming units (cfus) that maintained kanamycin resistance (and thus the target vector). All figures depicting these data show the log-transformed proportion of Kan^R^/total cfu, both with and without SpyCas9 induction. The growth curves in Supplemental Figure 1 match the experiment depicted Figure 1C for the uninduced SpyCas9 samples. For the uninduced *orf_1* sample, doxycycline was omitted from media throughout the experiment. Growth rates quoted in text were calculated using the slope of the OD600 growth curves during log phase, following a natural log transformation.

### Impact of AcrIIA22 on GFP expression

We swapped *spyCas9* for *egfp* in our CloDF13-based plasmid and co-expressed AcrIIA22 to determine if AcrIIA22 impacted expression from this construct. We reasoned that if AcrIIA22 influenced CloDF13’s copy number or the transcription of *spyCas9* it would also impact GFP levels in this construct (pCloDF13_GFP; Supplemental Table 4). To perform this experiment, we co-transformed pCloDF13_GFP and pZE21_tetR encoding *acrIIA22* into *E. coli* Turbo. Single colonies were picked into 4mL of LB containing spectinomycin 50 µg/ml (‘spec50’) and kanamycin 50 µg/ml (‘kan50’) and 0.5mM IPTG and grown overnight at 37°C shaking at 220rpm. The next morning the overnight culture was diluted 1:50 into both LB spec50 Kan50 + 0.5mM IPTG with and without doxycycline (to induce *acrIIA22*) and grown at 37°C for about 1.5 hours to mid-log phase (OD600 0.2-0.6). The OD600 was measured, and all samples were diluted to OD600 of 0.1 in two media types: (a) LB spec50 + kan50 + 0.5mM IPTG + 0.2% arabinose (inducing *gfp* only) or (b) LB spec50 + kan50 + 0.5mM IPTG + 0.2% arabinose + 100ng/ml doxycycline (inducing *gfp* and *acrIIA22*). A volume of 200 µl of each sample was then transferred to a 96-well plate in triplicate and we measured GFP fluorescence every 15 minutes for 24 hours (GFP was excited using 485 nm light and emission detected via absorbance at 528 nm). In parallel, we included control samples that lacked the kanamycin-marked plasmid and varied whether doxycycline was added or not (at 100 ng/ml). In these control samples, we noticed that doxycycline slightly diminished GFP expression (sub-toxic levels of the antibiotic may still depress translation). Thus, we normalized GFP fluoresced measurements in our experiment with AcrIIA22 to account for this effect in all +doxycycline samples. These fluorescence measurements are depicted in Supplemental Figure 2B.

### Western blots to AcrIIA22’s impact on SpyCas9 expression

Overnight cultures of *E. coli* Turbo that expressed pSpyCa9_crNT and pZE21_tetR encoding a gene of interest (Supplemental Tables 4, 5) were grown in LB spec50 + kan50 + 0.5mM IPTG. The next morning, these cultures were diluted 1:100 in 4ml of either (a) LB spec50 + kan50 + 0.5mM IPTG or (b) LB spec50 + kan50 + 0.5mM IPTG + 100 ng/ml doxycycline (to induce the gene of interest). We included samples that expressed either *acrIIA22* or *gfp* as a gene of interest. In all SpyCas9 constructs, we used a crRNA that did not target our plasmid backbone (pSpyCa9_crNT) to ensure that *acrIIA22* expression remained high and its potential impact on SpyCas9 expression levels would be most evident. All samples were grown for two hours at 37°C to reach mid-log phase (OD600 0.3 to 0.5) and transferred into media that contained 0.2% arabinose to induce SpyCas9. At transfer, volumes were normalized by OD600 value to ensure an equal number of cells were used (diluted to a final OD600 of 0.05 in the arabinose-containing medium). This second medium did or did not contain 100 ng/ml doxycycline to control expression of *acrIIA22* or *gfp,* as with the initial media. Throughout this experiment, we included a control strain that lacked pZE21_tetR and thus only expressed SpyCas9. Kanamycin and doxycycline were omitted from its growth media. For this control strain, we also toggled the addition of arabinose in the second growth medium to ensure positive and negative controls for SpyCas9 were included in our experiment. After three hours and six hours of SpyCas9 induction, OD600 readings were again taken and these values used to harvest an equal number of cells per sample (at three hours, OD600 values were between 0.76 and 0.93 and 0.75ml to 0.9ml volumes harvested; at six hours 0.4ml was uniformly harvested as all absorbance readings were approximately 1.6).

All samples were centrifuged at 4100g to pellet cells, resuspended in 100 µl of denaturing lysis buffer (12.5 mM Tris-HCl, pH 6.8; 4% SDS), and passed through a 25 gauge needle several times to disrupt the lysate. Samples were then boiled at 100°C for 10 minutes, spun at 13,000 rpm at 4°C for 15 minutes and the supernatants removed and frozen at −20°C. The next day, 12 µl of lysate was mixed with 4 µl of 4x sample buffer (200 mM Tris-HCl, 8% SDS, 40% glycerol, 200 mM DTT, and 0.05% bromophenol blue) and boiled at 100°C for 10 minutes. Then, 10 µl sample was loaded onto a BioRad Mini-Protean “any KD Stain Free TGX” gel (cat. #4569035) and run at 150V for 62 minutes. To verify that equivalent amounts of each sample were run, gels were visualized on a BioRad chemidoc for total protein content. Protein was then transferred to a 0.2 µM nitrocellulose membrane using the Bio-Rad Trans-Blot Turbo system (25 V, 1.3 A for 10 min). We then washed membranes in PBS/0.1% Triton-X before incubating them with a mixture of the following two primary antibodies, diluted in in Licor Odyssey Blocking Solution (cat. #927–40000): (i) monoclonal anti-SpyCas9, Diagenode cat. #C15200229-50, diluted 1:5,000; (ii) polyclonal anti-GAPDH, GeneTex cat. # GTX100118, diluted 1:5,000. The GAPDH antibody served as a second check to ensure equal protein levels were run. Membranes were left shaking overnight at 4°C, protected from light. Then, membranes were washed four times in PBS/0.1% Triton-X (ten-minute washes) before they were incubated for 30 minutes at room temperature with a mixture of secondary antibodies conjugated to infrared dyes. Both antibodies were diluted 1:15,000 in LiCor Odyssey Blocking Solution. To detect SpyCas9, the following secondary antibody was used: IR800 donkey, anti-mouse IgG, Licor cat# 926–32212. To detect GAPDH, IR680 goat, anti-rabbit IgG, Licor cat# 926-68071 was used. Blots were imaged on a Licor Odyssey CLx after three additional washes.

### Phage plaquing assay

Overnight cultures of *E. coli* Turbo that expressed pSpyCa9_crMu and pZE21_tetR encoding a gene of interest (Supplemental Tables 4, 5) were grown at 37°C in LB spec50 + kan50 + 0.5 mM IPTG. Genes of interest were either *acrIIA4*, *gfp*, or *acrIIA22*. The pSpyCas9 construct targeted phage Mu and was previously demonstrated to confer strong anti-phage immunity in this system^12^. A control strain expressing pZE21-tetR-*gfp* and SpyCas9_crNT (which encoded a CRISPR RNA that does not target phage Mu) was grown similarly. The next morning, all cultures were diluted 50-fold into LB spec50 + kan50 + 0.5 mM IPTG + 5 mM MgCl2 and grown at 37°C for three hours. Then, doxycycline was added to a final concentration of 100 ng/ml to induce the gene of interest. Two hours later, SpyCas9 was induced by adding a final concentration of 0.2% w/v arabinose. Two hours after that, cultures were used in soft-agar overlays on one of two media types, discordant for arabinose, to either maintain SpyCas9 expression or let it fade as arabinose was diluted in top agar and consumed by the host bacteria (per Supplemental Figure S2). Top and bottom agar media were made with LB spec50 + kan50 + 0.5 mM IPTG + 5 mM MgCl2. In cases where SpyCas9 expression was maintained, arabinose was also added at a final concentration of 0.02% to both agar types. Top agar was made using 0.5% Difco agar and bottom agar used a 1% agar concentration. For the plaquing assay, 100 µl of bacterial culture was mixed with 3 ml of top agar, allowed to solidify, and ten-fold serial dilutions of phage Mu spotted on top using 2.5 µl droplets. After the droplets dried, plates were overturned and incubated at 37°C overnight before plaques were imaged the subsequent day.

### Identification of AcrIIA22 homologs and hypervariable genomic islands

We searched for AcrIIA22 homologs in three databases: NCBI nr, IMG/VR, and a set of assembled contigs from 9,428 diverse human microbiome samples^18^. Accession numbers for the NCBI homologs are indicated on the phylogenetic tree in Figure 3A. They were retrieved via five rounds of an iterative PSI-BLAST search against NCBI nr performed on October 2^nd^, 2017. In each round of searching, at least 90% of the query protein (the original AcrIIA22 hit) was covered, 88% of the subject protein was covered, and the minimum amino acid identity of an alignment was 23% (minimum 47% positive residues; e-value ≤ 0.001). Only one unique AcrIIA22 homolog was identified in IMG/VR (from several different phage genomes) via a blastp search against the July, 2018 IMG/VR proteins database (using default parameters). It is identical to the sequence of AcrIIA22b (Figure 3A).

Most unique AcrIIA22 homologs were identified in the assembly data of over 9,400 human microbiomes performed by Pasolli and colleagues^18^. These data are grouped into multiple datasets: (i) the raw assembly data, and (ii) a set of unique species genome bins (SGBs), which was generated by first assigning species-level phylogenetic labels to each assembly and then selecting one representative genome assembly per species. We identified AcrIIA22 homologs using several queries against both databases. First, we performed a tblastn search against the SGB database using the AcrIIA22 sequence as a query, retrieving 141 hits from 137 contigs. A manual inspection of the genome neighborhoods for these hits revealed that most homologs originated from a short, hypervariable genomic island but that some homologs were encoded by prophages. No phage-finding software was used to identify prophages; they were apparent from a manual inspection of the gene annotations that neighbored *acrIIA22* homologs (see the section entitled “Annotation and phylogenetic assignment of metagenomic assemblies” for details).

To find additional examples of AcrIIA22 homologs and of these genomic islands, we then queried the full raw assembly dataset. To do so without biasing for *acrIIA22*-encoding sequences, we used the *purF* gene that flanked *acrIIA22*-encoding genomic islands as our initial query sequence (specifically, we used the *purF* gene from contig number 1 in Supplemental Table 3; its sequence is also in Supplemental Table 5). To consider only the recent evolutionary history of this locus, we required all hits have ≥98% nucleotide identity and required all hits to be larger than 15 kilobases in length to ensure sufficient syntenic information. From these contigs, we further filtered for those that had ≥98% nucleotide identity to *radC*, the gene which flanked the other end of *acrIIA22*-encoding genomic islands (again, we used the variant from contig number 1 in Supplemental Table 3; its sequence is also in Supplemental Table 5). In total, this search yielded 258 contig sequences; nucleotide sequences and annotations for these contigs are provided in Supplementary Dataset 5. We then searched for *acrIIA22* homologs in these sequences using tblastn, again observing them in genomic islands and prophage genomes (these prophages were assembled as part of the 258 contigs). In total, this search revealed 320 *acrIIA22* homologs from 258 contigs. The 258 genomic islands from these sequences were retrieved manually by extracting all nucleotides between the *purF* and *radC* genes. These extracted sequences were then clustered at 100% nucleotide identity with the sequence analysis software geneious to identify 128 unique genomic islands.

Combined, our two searches yielded 461 AcrIIA22 sequences from these metagenomic databases that spanned 410 contig sequences. The 461 AcrIIA22 homologs broke down into 410 that clustered with the genomic island-like sequences (we specifically searched for genomic islands) and 51 that clustered with prophage-like homologs (we never directly searched for prophages). We then combined these 461 AcrIIA22 sequences with those from NCBI and IMG/VR and clustered the group on 100% amino acid identity to reveal 30 unique proteins. To achieve this, we used the software cd-hit^43^ with the following parameters: -d 0 -g 1 -aS 1.0 -c 1.0. These 30 sequences were numbered to match their parent contig (as indicated in Supplemental Table 3) and used to create the phylogenetic tree depicted in Figure 3A. For AcrIIA22 homologs found outside NCBI, the nucleotide sequences and annotations their parent contigs can be found in Supplementary Datasets 1 and 2. This information can be retrieved for NCBI sequences via their accession numbers (which are shown in Figure 3A). The NCBI gene sequences also used in functional assays (Figure 3B) have been reprinted in Supplemental Table 5, for convenience.

### Annotation and phylogenetic assignment of metagenomic assemblies

Contig sequences from IMG/VR, the Pasolli metagenomic assemblies, and some NCBI entries lacked annotations, making it difficult to make inferences about *acrIIA22’s* genomic neighborhood. To facilitate these insights, we annotated all contigs as follows. We used the gene-finder MetaGeneMark^44^ to predict open reading frames (ORFs) using default parameters. We then used their amino acid sequences in a profile HMM search with HMMER3^45^ against TIGRFAM^46^ and Pfam^47^ profile HMM databases. The highest scoring profile was used to annotate each ORF. We annotated these contigs to facilitate genomic neighborhood analyses for *acrIIA22* and not to provide highly accurate functional predictions of their genes. Thus, we erred on the side of promiscuously assigning gene function and our annotations should be treated with the appropriate caution. From these annotated contigs, we immediately observed several examples of *acrIIA22*-encoding prophages (we noticed 35-40 kilobase insertions within some contigs that contained mostly co-linear genes with key phage functions annotated). As a simple means to sample this phage diversity, we manually extracted nine examples of these prophage sequences (their raw sequences and annotated genomes can be found in Supplementary Datasets 3 and 4). Annotations were imported to in the sequence analysis suite Geneious Prime 2020 v1.1 for manual inspection of genome neighborhoods.

We used the genome taxonomy database (GTDB) convention for all sequences discussed in this manuscript^48^. In part, this was because all *acrIIA22* genomes are found in *Clostridial* genomes, which are notoriously polyphyletic in NCBI taxonomies (for instance, the NCBI genus appears in GTDB genera and 29 GTDB families)^49^. All SGBs that we retrieved from the Pasolli assemblies were assigned taxonomy as part of that work and were called Clostridium sp. CAG-217. Similarly, NCBI assemblies that encoded the most closely *acrIIA22* homologs to our original hit were assigned to the GTDB genus CAG-217^48,49^. The raw assembly data from the Pasolli database was not assigned a taxonomic label but was nearly identical in nucleotide composition to the CAG-217 contigs (Figure 2, Supplementary Datasets 1 and 2). Therefore, we also refer to these sequences as originating in CAG-217 genomes but take care to indicate which sequences have been assigned a rigorous taxonomy and which ones for which taxonomy has been inferred in this fashion (Supplemental Table 3).

### Comparing genes in genomic islands to phage genomes

We first examined the annotated genes within each of the 128 unique genomic islands. Manual inspection revealed 54 unique gene arrangements (which differed in gene content and orientation). We then selected one representative from each arrangement and extracted all amino acid sequences from each encoded gene (n=506). Next, we collapsed these 506 proteins into orthologous groups by clustering at 65% amino acid using cd-hit with the following parameters: - d 0 -g 1 -aS 0.95 -c 0.65. These cluster counts were used to generate the histogram depicted in Figure 2C. To determine which protein families may also be phage encoded, the longest representative from each cluster with at least two sequences was queried against the database of nine CAG-217 phages described in the section entitled “Annotation and phylogenetic assignment of metagenomic assemblies”. We used tblastn with default parameters to perform this search, which revealed that some proteins in the CAG-217 genomic islands have homologs in prophage genomes that are out-of-frame with respect to the MetaGeneMark annotations depicted in Figure 2A.

### Phylogenetic tree of AcrIIA22 homologs

The 30 unique AcrIIA22 homologs we retrieved were used to create the phylogeny depicted in Figure 3A. These sequences were aligned using the sequence alignment tool in the sequence analysis suite Geneious Prime 2020 v1.1. This alignment is provided as Supplementary Dataset 6. From this alignment, the phylogenetic tree in Figure 3A was generated using PhyML with the LG substitution model and 100 bootstraps. Coloration and tip annotations were then added in Adobe Illustrator.

### Identification of CRISPR-Cas systems and Acrs in CAG-217 assemblies

To determine the type and distribution of CRISPR-Cas systems and Acrs in CAG-217 genomes, we downloaded all assembly data for the 779 SGBs assigned to CAG-217 in Pasolli *et. al*^18^ (bin 4303). We then predicted CRISPR-Cas systems for all 779 assemblies in bulk using the command line version of the CRISPR-Cas prediction suite, cctyper^50^. Specifically, we used version 1.2.1 of cctyper with the following options: --prodigal meta --keep_tmp. To identify type II-A Acrs, we first downloaded representative sequences for each of the 21 experimentally confirmed type II-A Acrs from the unified resource for tracking anti-CRISPRs^51^. We then used tblastn to query these proteins against the 779 CAG-217 genome bins and considered any hit with e-value better than 0.001 (which included all hits with >30% identity across 50% of the query). To check if these Acrs were present in *acrIIA22*-encoding phages, we performed an identical tblastn search, but this time used the set of nine *acrIIA22*-encoding prophages as a database.

### Recombinant protein overexpression and purification

The AcrIIA22 protein and its mutants were codon optimized for *E. coli* (Genscript or SynBio Technologies) and the gene construct was cloned into the pET15HE^12^ plasmid to contain an N-terminal, thrombin-cleavable 6XHistidine tag. Constructs were transformed and overexpressed in BL21 (DE3) RIL *E. coli* cells. A 10 mL overnight culture (grown in LB + 100 µg/mL ampicillin) was diluted 100-fold into the same media and grown at 37°C with shaking to an OD600 of 0.8, followed by induction with 0.5 mM IPTG. The culture was shaken for an additional 3 hours at 37°C. Cells were harvested by centrifugation and the pellet stored at −20°C until purification.

Cell pellets were resuspended in 25 mM Tris, pH 7.5, 300 mM NaCl, 20 mM imidazole (Lysis Buffer) and lysed by sonication on ice. The lysate was centrifuged in an SS34 rotor at 18,000 rpm for 25 minutes, followed by filtering through a 5 µm syringe filter (Millipore #SLSV025LS). The clarified lysate was bound using the batch method to Ni-NTA agarose resin (Qiagen) at 4°C for 1 hour. The resin was transferred to a gravity column (Biorad), washed with >50 column volumes of Lysis Buffer and eluted with 25 mM Tris, pH 7.5, 300 mM NaCl, 200 mM imidazole. The protein was diluted with 2 column volumes of 25 mM Tris, pH 7.5 and purified on a HiTrapQ column (GE Healthcare) using a 20 mL gradient from 150 mM to 1 M NaCl in 25 mM Tris, pH 7.5. Peak fractions were pooled, concentrated and buffer exchanged into 200 mM NaCl, 25 mM Tris, pH 7.5 using an Amicon Ultra centrifugal filter with a 3,000 molecular weight cutoff (Millipore, UFC900324), then cleaved in an overnight 4°C incubation with biotinylated thrombin (EMD Millipore). Streptavidin agarose slurry (Novagen) was incubated with cleaved protein at 4°C for 30 minutes to remove thrombin. The sample was then passed through a 0.22 µm centrifugal filter and loaded onto a HiLoad 16/60 Superdex 200 prep grade size exclusion column (Millipore Sigma) equilibrated in 25 mM Tris, pH 7.5, 200 mM NaCl. The peak fractions were confirmed for purity by SDS-PAGE. Figure 4C depicts size exclusion chromatography data generated for thrombin-cleaved AcrIIA22 variants generated using a Superdex75 16/60 (GE HealthCare) column with 25 mM Tris, pH 7.5, 200mM NaCl. Recombinant AcrIIA4 was purified similarly to other Acr proteins as previously described^12^, but with the following deviations. First, the 6XHistidine-tagged AcrIIA4 gene was cloned into pET15B rather than pET15HE, which differs by only by a few bases just upstream of the N-terminal thrombin tag. IPTG was used at 0.2 mM and cells were harvested after 18 hours of induction at 18°C. Thrombin cleavage also occurred at 18°C. This untagged version was used to help generate Supplemental Figure 5. Peak fractions for all proteins were pooled, concentrated, flash frozen as single-use aliquots in liquid nitrogen, and stored at −80°C.

SpyCas9 was expressed in E. coli from plasmid pMJ806 (addgene #39312) to contain a TEV-cleavable N-terminal 6XHis-MBP tag and was purified as described previously^12^. Briefly, sequential steps of purification consisted of Ni-NTA affinity chromatography, TEV cleavage, Heparin HiTrap chromatography and SEC. The protein was stored in a buffer consisting of 200 mM NaCl, 25 mM Tris (pH 7.5), 5% glycerol, and 2 mM DTT.

To perform *in vitro* pulldown experiments, we purified AcrIIA22 and AcrIIA4 proteins with a C-terminal twin-strep tag. To achieve this, the Acrs were subcloned into pET15B which was previously engineered to contain a thrombin-cleavable C-terminal twin-strep tag. The protein was expressed as described above and purified according to the manufacturer’s guidelines (IBA Inc.). Briefly, cell lysates were resuspended in Buffer W (150 mM NaCl, 100 mM Tris, pH 8.0, 1 mM EDTA) and lysed by sonication. Clarified lysates were then passed over Streptactin-Sepharose resin using a gravity filtration column. The flow through was passed over the resin an additional time. The column was washed with a minimum of 20 column volumes of buffer W, followed by elution in buffer E (150 mM NaCl, 100 mM Tris, pH 8.0 mM, 1 EDTA, 2.5 mM desthiobiotin). The eluted protein was purified over a HiTrap Q column, followed by SEC in 200 mM NaCl, 25 mM Tris, 7.5.

### X-ray crystallography and structural analyses

An AcrIIA22 crystal was grown using 14mg/mL protein via the hanging drop method using 200mM ammonium nitrate, 40% (+/-)-2-methyl-2,4-pentanediol (MPD, Hampton Research), 10mM MgCl2 as a mother liquor. Diffraction data was collected at the Argonne National Laboratory Structural Biology Center synchrotron facility (Beamline 19BM). Data was processed with HKL2000 in space group P4332, then built and refined using COOT^52^ and PHENIX^53^. The completed 2.80Å structure was submitted to the Protein Data Bank with PDB Code 7JTA. We submitted this finished coordinate file to the PDBe PISA server (Protein Data Bank Europe, Protein Interfaces, Surfaces and Assemblies; http://pdbe.org/pisa/) which uses free energy and interface contacts to calculate likely multimeric assemblies^25^. The server calculated tetrameric, dimeric and monomeric structures to be thermodynamically stable in solution. The tetrameric assembly matches the molecular weight expected from the size exclusion column elution peak and is the most likely quaternary structure as calculated by the PISA server. The tetramer gains −41.8 kcal/mol free energy by solvation when formed and requires an external driving force of 3.1 kcal/mol to disassemble it according to PISA ΔG calculations.

### sgRNA generation

The single-guide RNA (sgRNA) for use in *in-vitro* experiments was generated as described previously^12^. It was transcribed from a double-stranded DNA (dsDNA) template by T7 RNA polymerase using Megashortscript Kit (Thermo Fisher #AM1354). We made the dsDNA template via one round of thermal cycling (98°C for 90 s, 55°C for 15 s, 72°C for 60 s) in 50 µl reactions. We used the Phusion PCR polymerase mix (NEB) containing 25 pmol each of the following two oligo sequences (the sequence that binds the protospacer on our pIDTsmart target vector is underlined):

i. GAAATTAATACGACTCACTATAGG<u>TAATGAAATAAGATCACTAC</u>GTTTTAGAGCT AGAAATAGCAAGTTAAAATAAGGCTAGTCCG
ii. AAAAAAGCACCGACTCGGTGCCACTTTTTCAAGTTGATAACGGACTAGCCTTATTTTAACTTGC.

The dsDNA templates were then purified using an Oligo Clean and Concentrator Kit (ZymoResearch) before quantification via the Nanodrop. Reactions were then treated with DNAse, extracted via phenol-chloroform addition followed by chloroform, ethanol precipitated, resuspended in RNase free water, and frozen at −20°C. RNA was quantified by Nanodrop and analyzed for quality on 15% acrylamide/TBE/UREA gels.

### Pulldown assay using strep-tagged AcrIIA22 and AcrIIA4

The same buffer was used for pulldowns and to dilute proteins, consisting of 200 mM NaCl, 25 mM Tris (pH 7.5). As a precursor to these assays, 130 pmol SpyCas9 and sgRNA were incubated together at room temperature for 15 minutes where indicated. SpyCas9, with or without pre-complexed sgRNA, was then incubated with 230 pmol AcrIIA4 or 320 pmol AcrIIA22 for 25 minutes at room temperature. Subsequently, 50 µl of a 10% slurry of Streptactin Resin (IBA biosciences #2-1201-002), pre-equilibrated in binding buffer, was added to the binding reactions and incubated at 4°C on a nutator for 45 minutes. Thereafter all incubations and washes were carried out at 4°C or on ice. Four total washes of this resin were performed, which included one tube transfer. Washes proceeded via centrifugation at 2000 rpm for one minute, aspiration of the supernatant with a 25-gauge needle, and resuspension of the beads in 100 µl binding buffer. Strep-tagged proteins were eluted via suspension in 40 µl of 1x BXT buffer (100 mM Tris-Cl, 150 mM NaCl, 1 mM EDTA, 50 mM Biotin, pH 8.0) and incubated for 15 min at room temperature. After centrifugation, 30 µl of supernatant was removed and mixed with 4X reducing sample buffer (Thermo Fisher). Proteins then separated by SDS PAGE on BOLT 4–12% gels in MES buffer (Invitrogen) and visualized by Coomassie staining.

### SpyCas9 linear DNA cleavage assay

All SpyCas9 cleavage reactions using linear DNA were performed in the following cleavage buffer: 20mM Tris HCl (pH7.5), 5% glycerol, 100mM KCl, 5mM MgCl2, 1mM DTT. In preparation for these reactions, all proteins were diluted in 30 mM NaCl / 25 mM Tris, pH 7.4 / 2.7mM KCl, whereas all DNA and sgRNA reagents were diluted in nuclease-free water. Where indicated, SpyCas9 (0.36 µM) was incubated with sgRNA (0.36 µM) for 10 minutes at room temperature. Before use, sgRNA was melted at 95°C for five minutes and then slowly cooled at 0.1 °C/s to promote proper folding. SpyCas9 (either pre-complexed with sgRNA or not, as indicated in Supplemental Figure 6) was then incubated for 10 minutes at room temperature with AcrIIA4 (2.9 µM) or AcrIIA22 at the following concentrations: [23.2, 11.6, 5.8, and 2.9 µM]. As substrate, the plasmid pIDTsmart was linearized by restriction digest and used at a final concentration of 3.6 nM. The reaction was initiated by the addition of this DNA substrate in isolation or in combination with sgRNA (0.36 µM) as indicated in Supplemental Figure 6. Reactions were immediately moved to a 37°C incubator and the reaction stopped after fifteen minutes via the addition of 0.2% SDS/100 mM EDTA and incubating at 75°C for five minutes. Samples were then run on a 1.5% TAE agarose gel at 120V for 40 minutes. Densitometry was used to calculate the proportion of DNA cleaved by SpyCas9 via band intensities quantified using the BioRad ImageLab software v5.0.

### *In vivo* assay to assess impact of AcrIIA22 on plasmid topology

In all experiments, cultures were first grown overnight at 37°C with shaking at 220 rpm in LB with 0.5mM IPTG, spectinomycin (at 50 µg/mL), and kanamycin (at 50 µg/mL). Then, these overnight cultures were diluted 1:50 into LB with 0.5mM IPTG, spectinomycin (at 50 µg/mL), and, where indicated, doxycycline (at 100 ng/mL, to induce *acr*s). Cultures were grown at 37°C with shaking at 220 rpm and, if indicated, 0.2% (L)-arabinose was added after two hours of growth to induce *spyCas9* expression. The next morning, cultures were centrifuged at 4100*g* and plasmids purified using a miniprep kit (Qiagen). The concentration of dsDNA in each miniprep was measured using the Qubit-4 fluorometer and the associated dsDNA high sensitivity assay kit (Invitrogen). For each sample with a SpyCas9-expressing plasmid, 150ng of DNA was digested with the restriction enzyme HincII (NEB) per manufacturer’s recommendations, except that digests were incubated overnight before being stopped by heating at 65°C for 20 minutes. This restriction enzyme will cut once, only in the SpyCas9 plasmid, to linearize it. This allowed us to visualize the SpyCas9 plasmid as a single band, which served two purposes: (i) it allowed us to more easily identify bands from *acrIIA22*-encoding plasmids (which had not been digested), and (ii) it served as an internal control for plasmid DNA that is unaffected by SpyCas9 targeting or AcrIIA22 expression (Supplemental Figure 2). Following restriction digest, 30ng of sample was analyzed via gel electrophoresis using a 1% TAE-agarose gel run at 120V for between 45 and 60 minutes. In samples that lacked a SpyCas9-expressing plasmid, 30ng of purified plasmid was directly analyzed by gel electrophoresis, as described previously.

### *In vitro* AcrIIA22 plasmid nicking assay

Except for the divalent cation experiment, all reactions were performed using NEB buffer 3.1 (100 mM NaCl, 50 mM Tris-HCl, pH 7.9, 10 mM MgCl2, 100 µg/mL BSA). To determine cation preference, the same reaction buffer was re-created, but MgCl2 was omitted. All proteins were diluted in 130 mM NaCl, 25 mM Tris, pH 7.4, 2.7 mM KCl. DNA was diluted in nuclease-free water. In the cation preference experiment, 60 µM AcrIIA22 and 6 nM of purified pIDTsmart plasmid DNA were used. All other reactions were set up with the AcrIIA22 final concentrations indicated in Figure 5 and Supplemental Figure 7. In the cation preference experiment, reactions were started by adding 10 mM of the indicated cation. All other reactions were initiated via the addition of 2 nM pIDTsmart plasmid DNA. In all cases, reactions were immediately transferred to a 37°C incubator. At 0.5, 1, 2, 4, 6, or 20-hour timepoints, a subset of the reaction was removed and run on a 1% TAE agarose gel at 120V for 40 minutes. For the cation preference experiment, only the 2-hour timepoint was considered and the reaction was stopped via the addition of NEB loading buffer and 100 mM EDTA. In this case, DNA was visualized on a 1% TBE gel run for 60 minutes at 110V. Densitometry was used to calculate the proportion of DNA in each topological form via band intensities quantified using the BioRad ImageLab software v5.0.

### SpyCas9 cleavage kinetics assay

Except where indicated in Supplemental Figure 9B, all cleavage reactions were performed in the following cleavage buffer: 20mM Tris HCl (pH7.5), 5% glycerol, 100mM KCl, 5mM MgCl2, 1mM DTT. In preparation for these reactions, all proteins were diluted in 30 mM NaCl / 25 mM Tris, pH 7.4 / 2.7mM KCl, whereas all DNA and sgRNA reagents were diluted in nuclease-free water. NEB Buffer 3.1 (100 mM NaCl, 50 mM Tris-HCl, pH 7.9, 10 mM MgCl2, 100 µg/mL BSA) was used as a reaction buffer in Supplemental Figure 9B.

In preparation for these reactions, purified pIDTsmart plasmid was pre-treated with either AcrIIA22, the nickase Nb.Bss.SI (NEB), or no enzyme. For the AcrIIA22 pre-treatment, 3.1 µg of plasmid was incubated with 230 µM AcrIIA22 and the plasmid nicked as described previously. Plasmid nicking with Nb.Bss.SI proceeded via manufacturer’s recommendations (NEB). Both reactions were incubated at 37 °C for 2 hours. To isolate the nicked plasmid, samples were then run on a 1.5% agarose gel for 2 hours and the open-circle form of the plasmid was excised and purified using the Zymo Research Gel DNA Recovery Kit. Untreated plasmid was also purified via gel extraction. Plasmid yield was quantified using a Nanodrop.

To determine SpyCas9’s substrate preference, we incubated each pre-treated plasmid substrate with SpyCas9 and looked for the appearance of a linearized plasmid as indication of SpyCas9 digestion. In all cases, SpyCas9 was used at a final concentration of 31.2 nM. To begin the reaction, DNA substrate and sgRNA were added simultaneously to the reaction mix and the samples moved immediately from ice to 37 °C and incubated for either 1 or 5 minutes. We noticed that the digestion reaction proceeded too quickly with NEB Buffer 3.1 to detect SpyCas9’s substrate preference (i.e., the substrates were all rapidly linearized Supplemental Figure 9B). The cleavage buffer used in most reactions (detailed atop this section) was chosen because it slowed digestion kinetics so that we could detect SpyCas9’s substrate preference. Before addition to the reaction, sgRNA was melted at 95°C for five minutes and then slowly cooled at 0.1 °C/s to promote proper folding. At each timepoint, 5 µl of the reaction was removed and the reaction was stopped using 0.2% SDS/100 mM EDTA, then incubating at 75°C for 5 minutes. Samples were run on a 1.5% TAE gel at 120V for 40 minutes.

## Acknowledgements

We thank Kaylee Dillard, Ilya Finkelstein, and Tera Levin for comments on the manuscript. Use of the Advanced Photon Source, an Office of Science User Facility operated for the U.S. Department of Energy (DOE) Office of Science by Argonne National Laboratory, was supported by the U.S. DOE under Contract No. DE-AC02-06CH11357. This work was supported by a Helen Hay Whitney Foundation postdoctoral fellowship awarded to KJF, a Seattle University summer faculty fellowship to BKK, NIH grant R01GM105691 and discretionary funding from the Fred Hutchinson Cancer Research Center to BLS, and grants from the G. Harold and Leila Y. Mathers Foundation and the Howard Hughes Medical Institute to HSM. The funders played no role in study design, data collection and interpretation, or the decision to publish this study. HSM is an Investigator of the Howard Hughes Medical Institute.

---

**Supplemental Figure 1.**
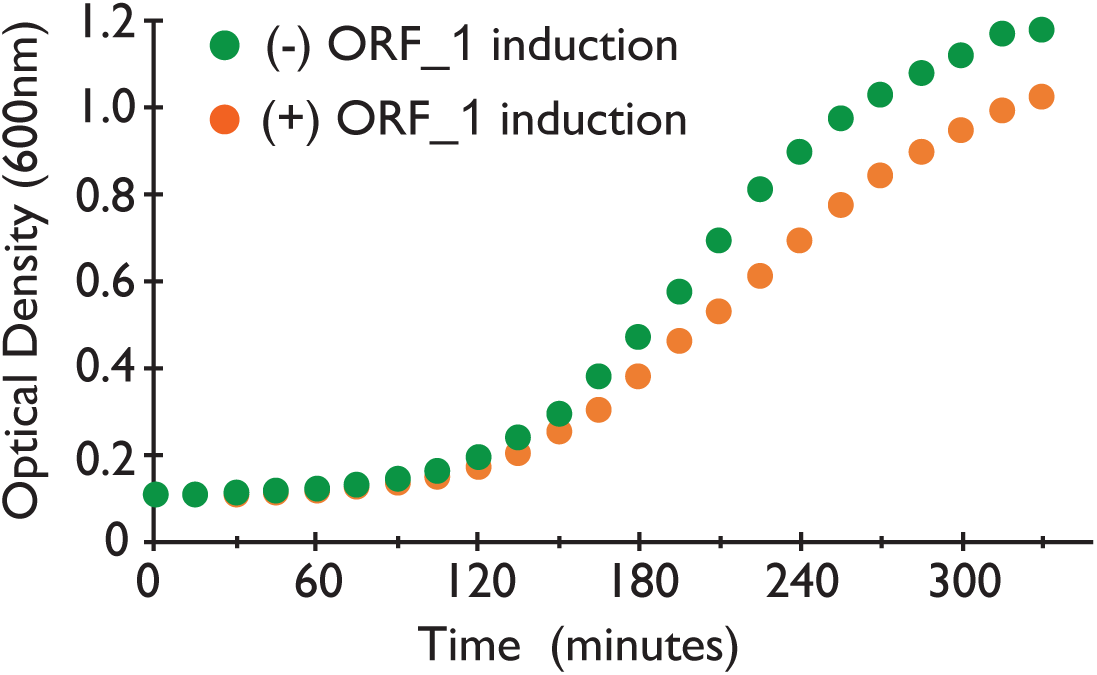
*Orf_1* confers mild toxicity in *E. coli*. Growth curves with (orange) and without (green) *orf_1* induction. These growth data map directly to the cfu data in Figure 1C, demonstrating that anti-SpyCas9 activity occurs under conditions with minimal *orf_1* toxicity. Samples were removed after six hours of growth to plate for cfus. Growth curves are shown for samples without SpyCas9 induction to ensure that *orf_1* toxicity is not mitigated due to elimination of its plasmid. Points indicate averages from three replicates. Standard deviations at each timepoint are so small that the error bars do not exceed the bounds of the data point.

**Supplemental Figure 2.**
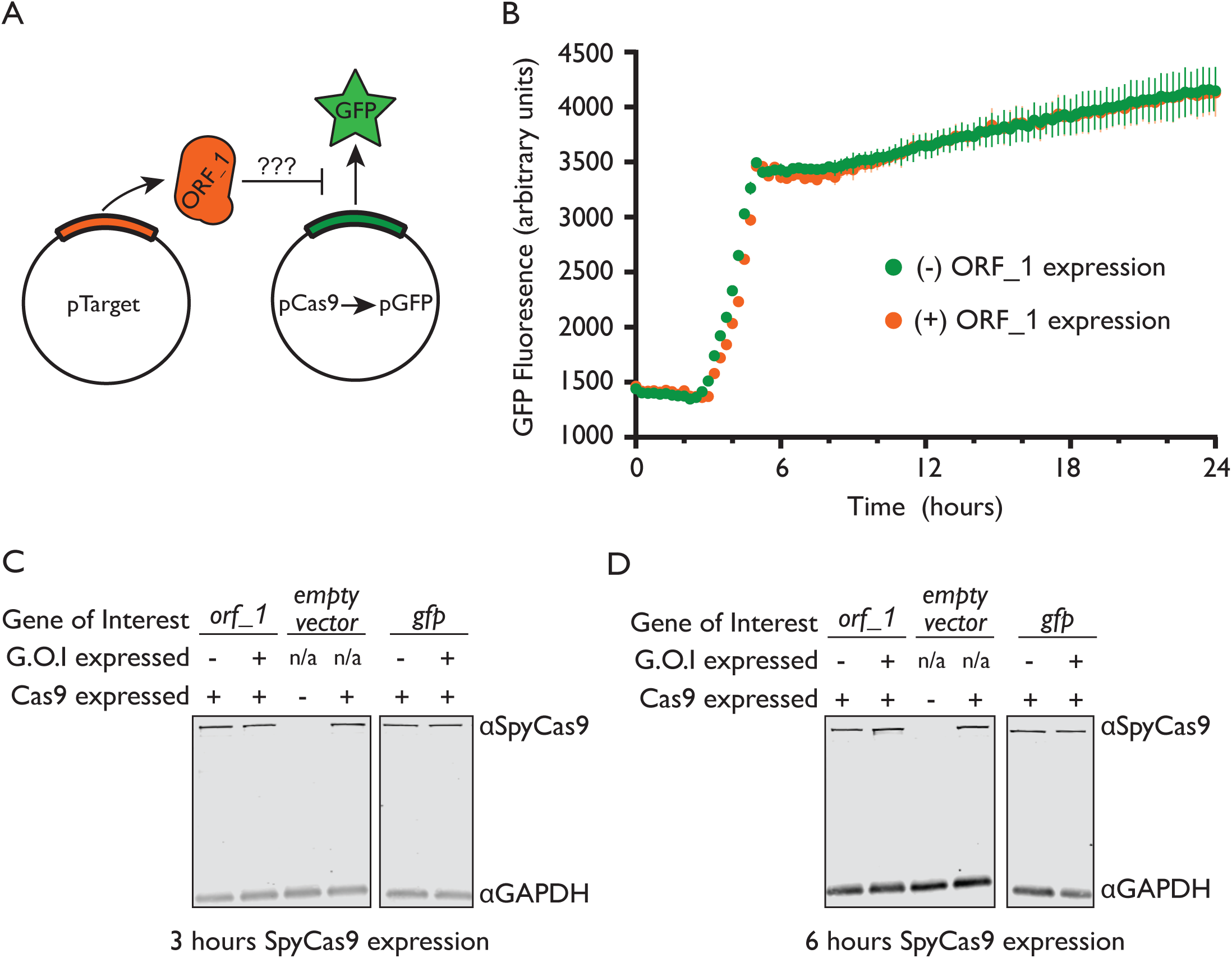
*Orf_1* does not impact SpyCas9 expression. (**A**) Cartoon depicting the experimental design for (**B**). If ORF_1 prevented transcription from pCas9 or altered its copy number, we would expect expression of the *orf_1* gene to deplete the level of green fluorescence observed from a construct that replaces the *spycas9* gene with *gfp*. (**B**) Fluorescence measurements for the experiment depicted in panel A, throughout an *E. coli* growth curve. ORF_1 does not impact GFP expression. Points indicate averages from three replicates, error bars indicate standard deviation. (**C**) A western blot to detect SpyCas9 expression as a function of ORF_1 or GFP expression in growing *E. coli* cultures. As an internal control, GAPDH expression was also detected. (**D**) As in panel C, but samples were collected six hours after SpyCas9 induction, instead of three.

**Supplemental Figure 3.**
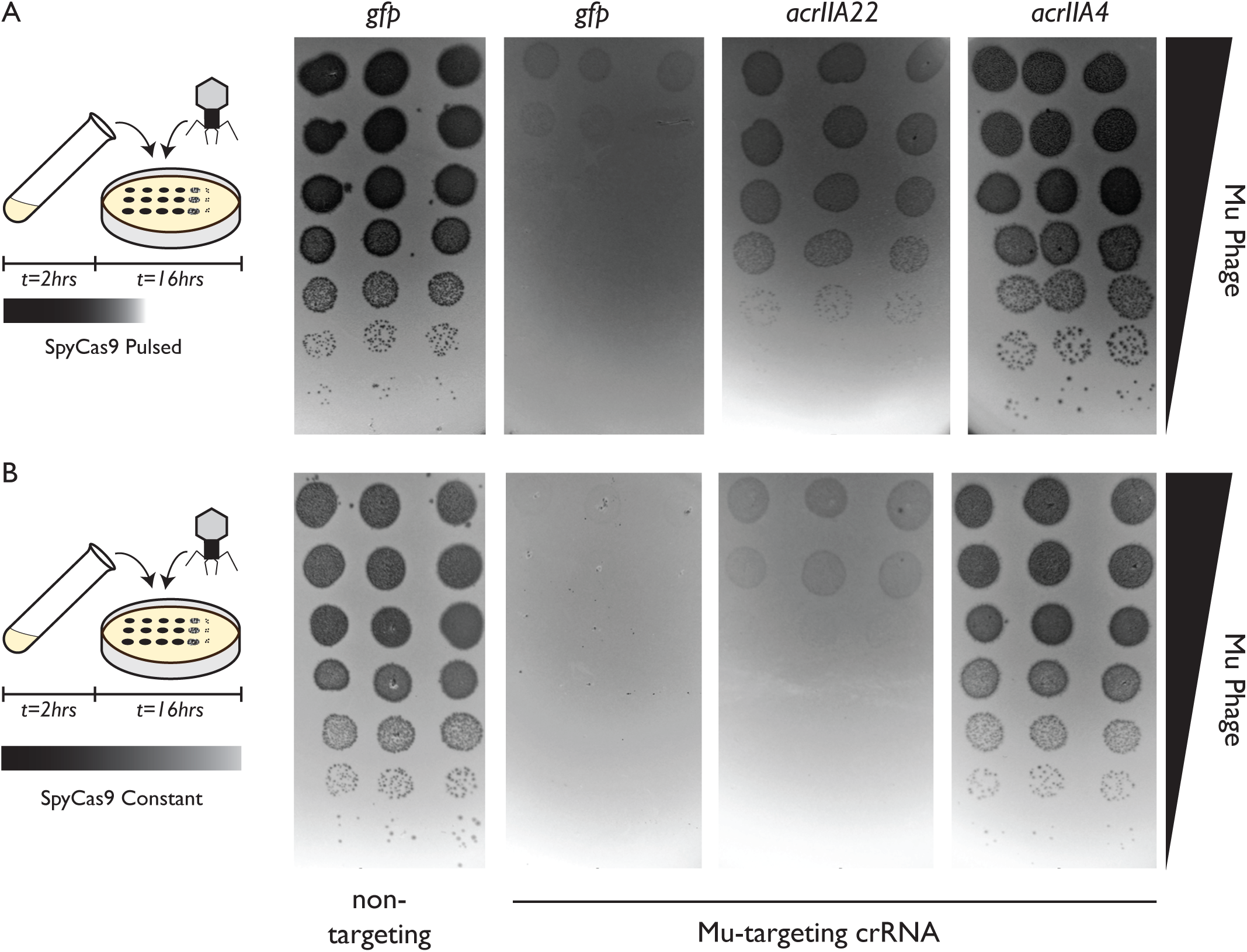
Mu phage fitness, measured by plaquing on *E. coli*. Plaquing is measured in the presence of *gfp*, *acrIIA22*, or *acrIIA4* via serial ten-fold dilutions. Bacterial clearing (black) occurs when phage Mu overcomes SpyCas9 immunity and lyses *E. coli*. In (**A**) and in (**B**), SpyCas9 with a Mu-targeting crRNA confers substantial protection against phage Mu relative to a non-targeting (n.t.) control, in both conditions tested. These conditions are depicted at left, with the only difference being whether SpyCas9 was only expressed in liquid growth prior to phage infection (panel A) or expressed both in liquid media and in solid media throughout infection (panel B). The positive control *acrIIA4* significantly enhances Mu fitness by inhibiting SpyCas9 in all conditions. In contrast, *acrIIA22* confers milder protection against SpyCas9. The indicated *acr* gene or *gfp* control is expressed from a second plasmid, in trans.

**Supplemental Figure 4.**
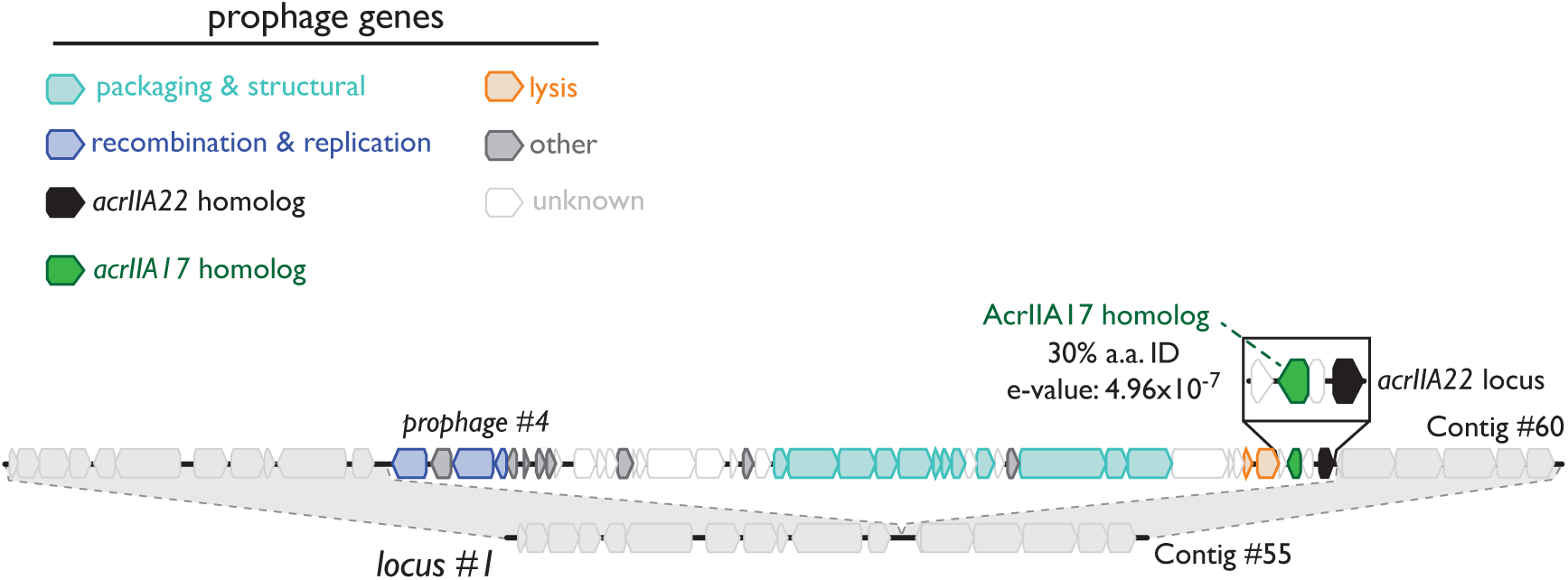
An *acrIIA22*-encoding prophage similar to those depicted in Figure 2A. Prophage genes are colored by functional category, per the legend and as in Figure 2. This prophage encodes for a homolog of the previously described SpyCas9 inhibitor *acrIIA17* within one kilobase of an *acrIIA22* homolog. Sequence relatedness for the depicted *acrIIA17* gene and the original discovery by Mahendra *et al*. is shown. Because phages often encode multiple *acr*s in the same locus, the co-localization of *acrIIA17* with *acrIIA22* is consistent with the latter gene functioning natively to inhibit CRISPR-Cas activity. Contigs are numbered to indicate their descriptions in Supplemental Table 3, which contains their metadata, taxonomy, and sequence retrieval information. All sequences and annotations may also be found in Supplementary Datasets 1 and 2.

**Supplemental Figure 5.**
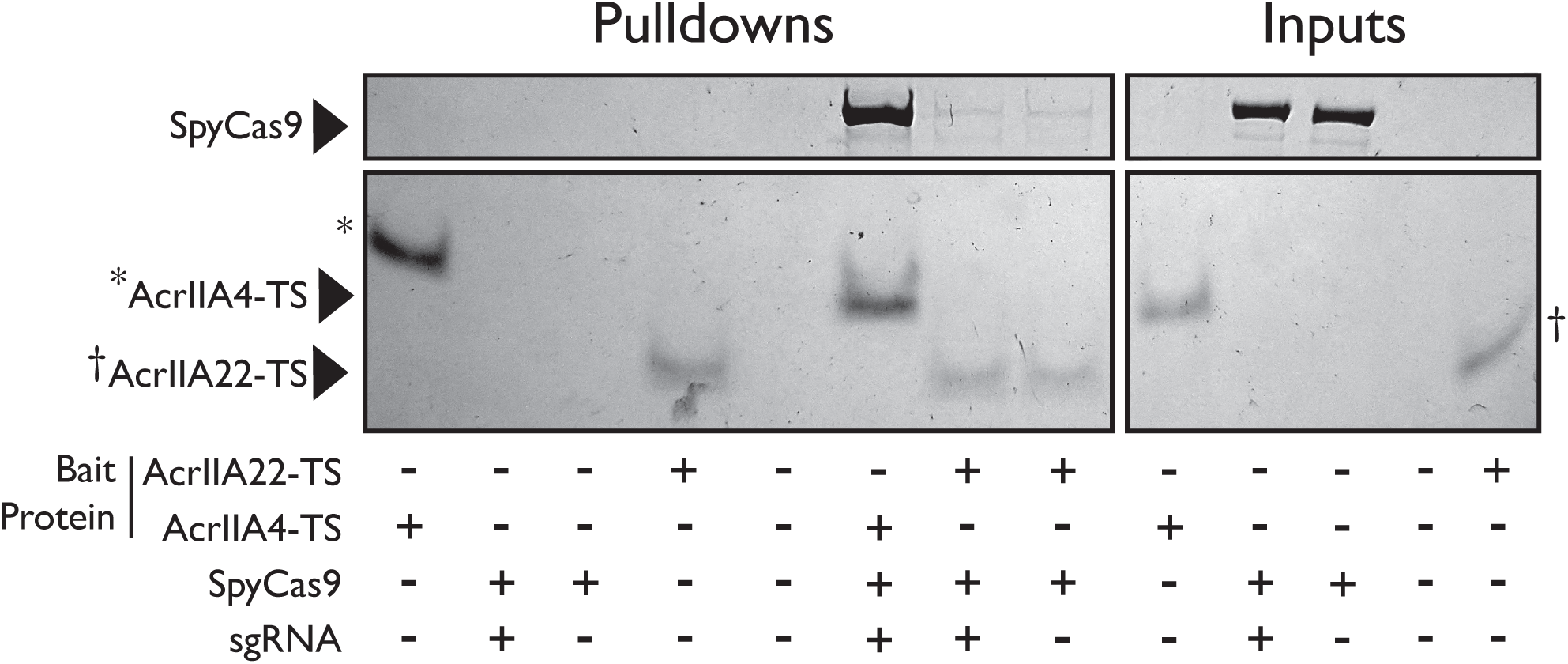
AcrIIA22 does not strongly bind SpyCas9. SpyCas9 and sgRNA were pre-incubated before mixing with a twin-strep (TS) tagged AcrIIA22 or AcrIIA4. SpyCas9 without sgRNA was also used. (A) Streptactin pulldowns on AcrIIA4 also pulled down SpyCas9 pre-incubated with sgRNA, as previously reported. Similar pulldowns with AcrIIA22 indicate little to no interaction with SpyCas9, regardless of whether sgRNA was used. These images depict total protein content visualized by Coomassie stain. Reaction components are indicated below the gel image. Aterisks (*) and dagger (†) symbols indicate AcrIIA4 and AcrIIA22 protein bands that run at slightly different positions than expected due to gel smiling.

**Supplemental Figure 6.**
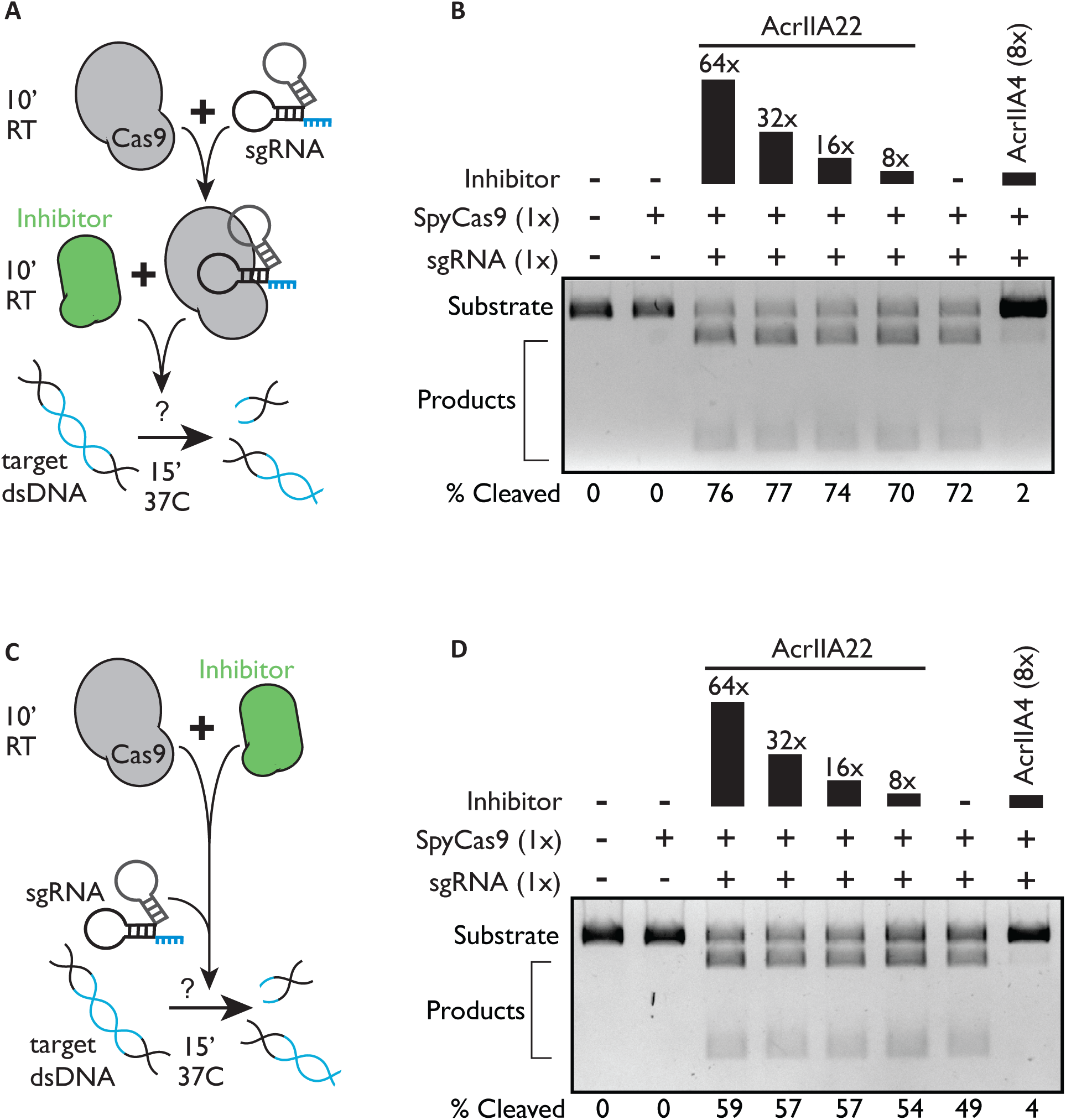
AcrIIA22 does not inhibit SpyCas9 from cleaving linear DNA. (**A**) Cartoon depicting the experiment in (**B**). SpyCas9 was pre-incubated with sgRNA targeting linear DNA. Then, Acrs candidates were added. Subsequently, cleavage reactions were performed, and the DNA products visualized by gel electrophoresis in panel B. (**B**) Products of the reactions described in panel A for the inhibitors AcrIIA22 and AcrIIA4. Reaction components are depicted atop the gel image, with molar equivalents relative to SpyCas9 indicated. The percent of DNA substrate cleaved by SpyCas9 is quantified below each lane. (**C**) As in panel A, except candidate Acrs were incubated with SpyCas9 before sgRNA addition. Reactions were begun via the simultaneous addition of sgRNA and linear dsDNA instead of dsDNA in isolation. (**D**) The products of the reactions described in panel C for AcrIIA22 and AcrIIA4 inhibitors. To push these Cas9 digestion reactions toward completion, ratios of Cas9:DNA were ten-fold higher than those shown in Figure 6C and reactions were allowed to progress for three times as long.

**Supplemental Figure 7.**
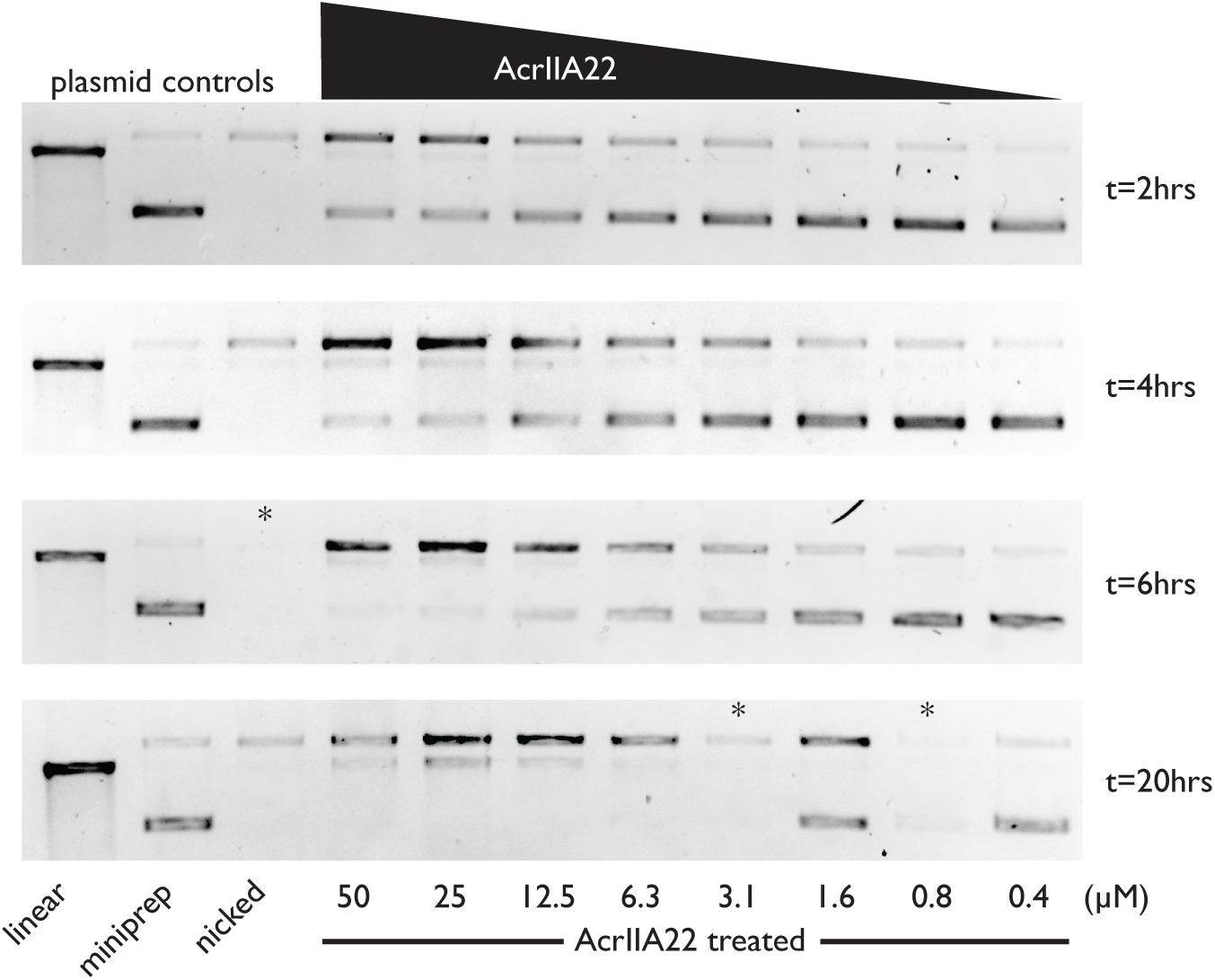
AcrIIA22 nicks supercoiled plasmids *in vitro*. Plasmid controls are in the leftmost three lanes. Reaction times are indicated to the right of each gel. AcrIIA22 nicks supercoiled plasmids in a concentration and time dependent manner. Extended incubations at high concentrations produce a linearized product. Asterisks (*) indicate loading errors, where less sample was loaded than other lanes.

**Supplemental Figure 8.**
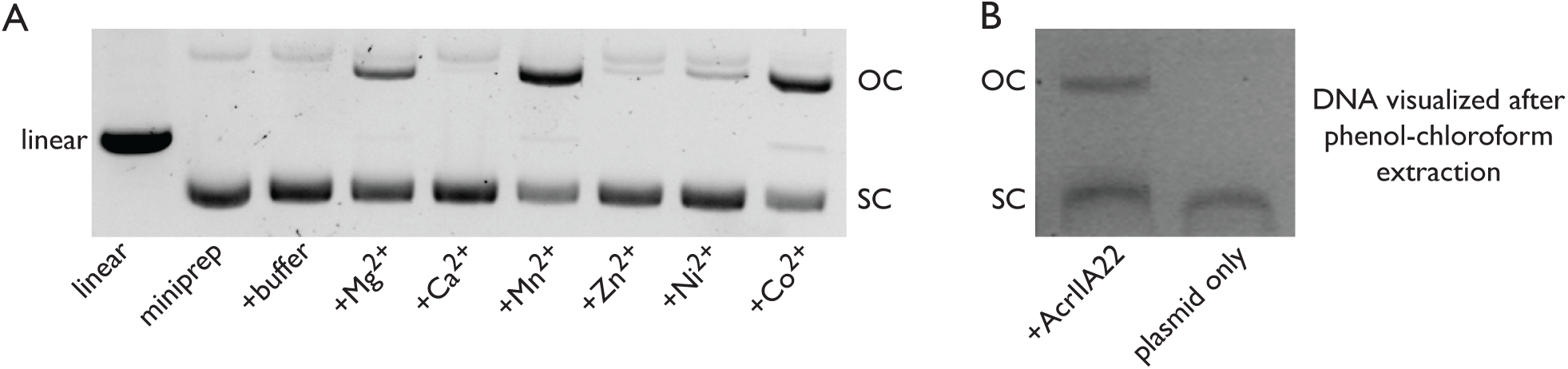
AcrIIA22 is a nickase. (**A**) The impact of different divalent cations on AcrIIA22’s nickase activity. OC, open-circle plasmid form. SC, supercoiled plasmid. (**B**) The open-circle plasmid product persists through phenol-chloroform extraction following AcrIIA22 treatment.

**Supplemental Figure 9.**
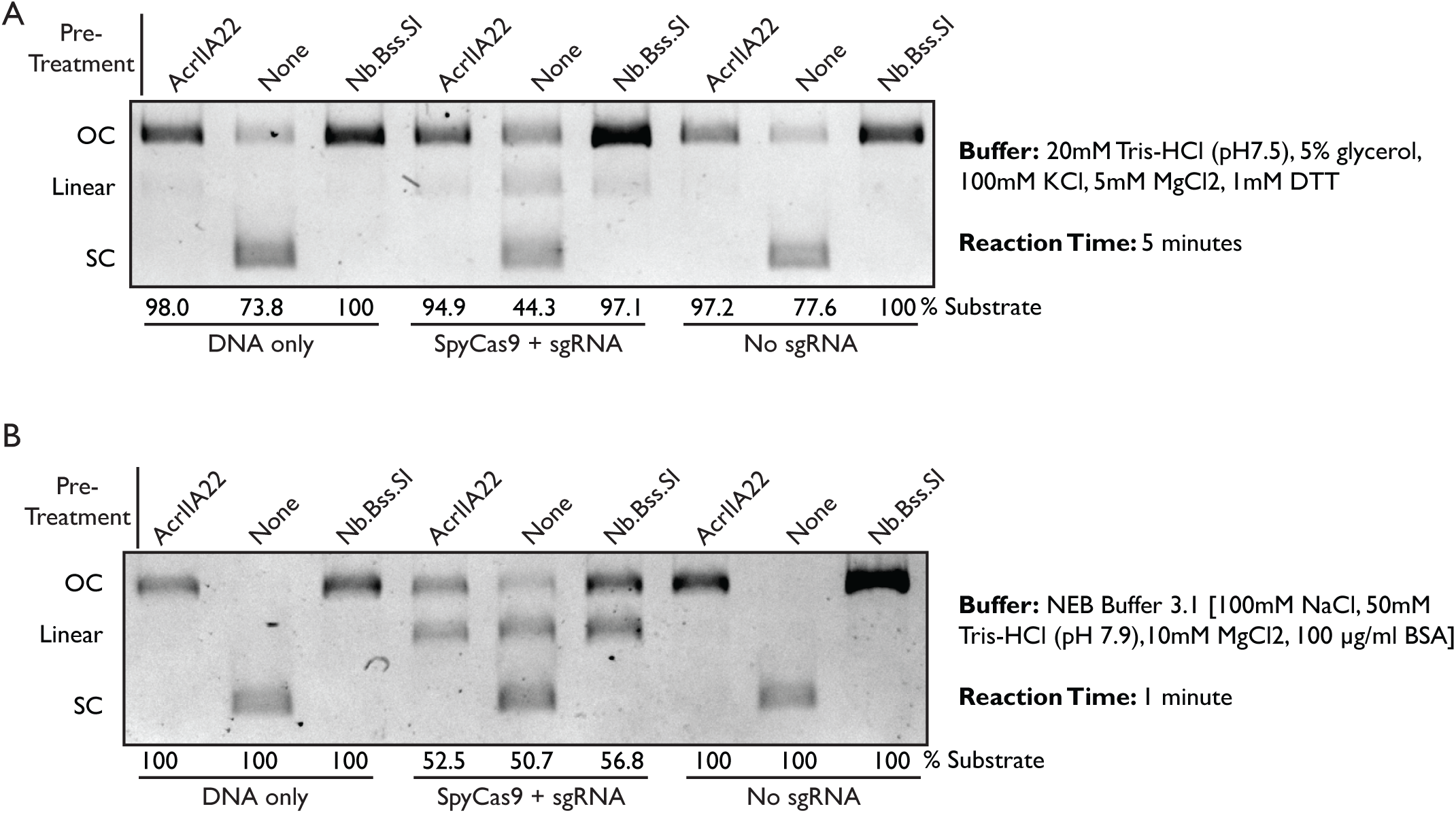
Nicked plasmids are less susceptible to linearization via SpyCas9 cleavage. (**A**) As in Figure 6C, Plasmid purifications from overnight cultures were either left unmodified or pre-treated with one of two nickase enzymes, AcrIIA22 or Nb.Bss.SI. Linear, open-circle (OC), and supercoiled (SC) plasmid forms are indicated. The % substrate value indicates the percentage of DNA in the nicked form for AcrIIA22 or Nb.Bss.SI-treated plasmids or in the supercoiled form for the untreated miniprep. Reaction components are listed below each lane. Buffer conditions and reaction time is listed at right. (**B**) As in (**A**), but with different reaction conditions, listed at right. In these conditions, the reaction proceeded too quickly to detect SpyCas9’s substrate preference (all substrates were rapidly linearized).

**Supplemental Table 1.**
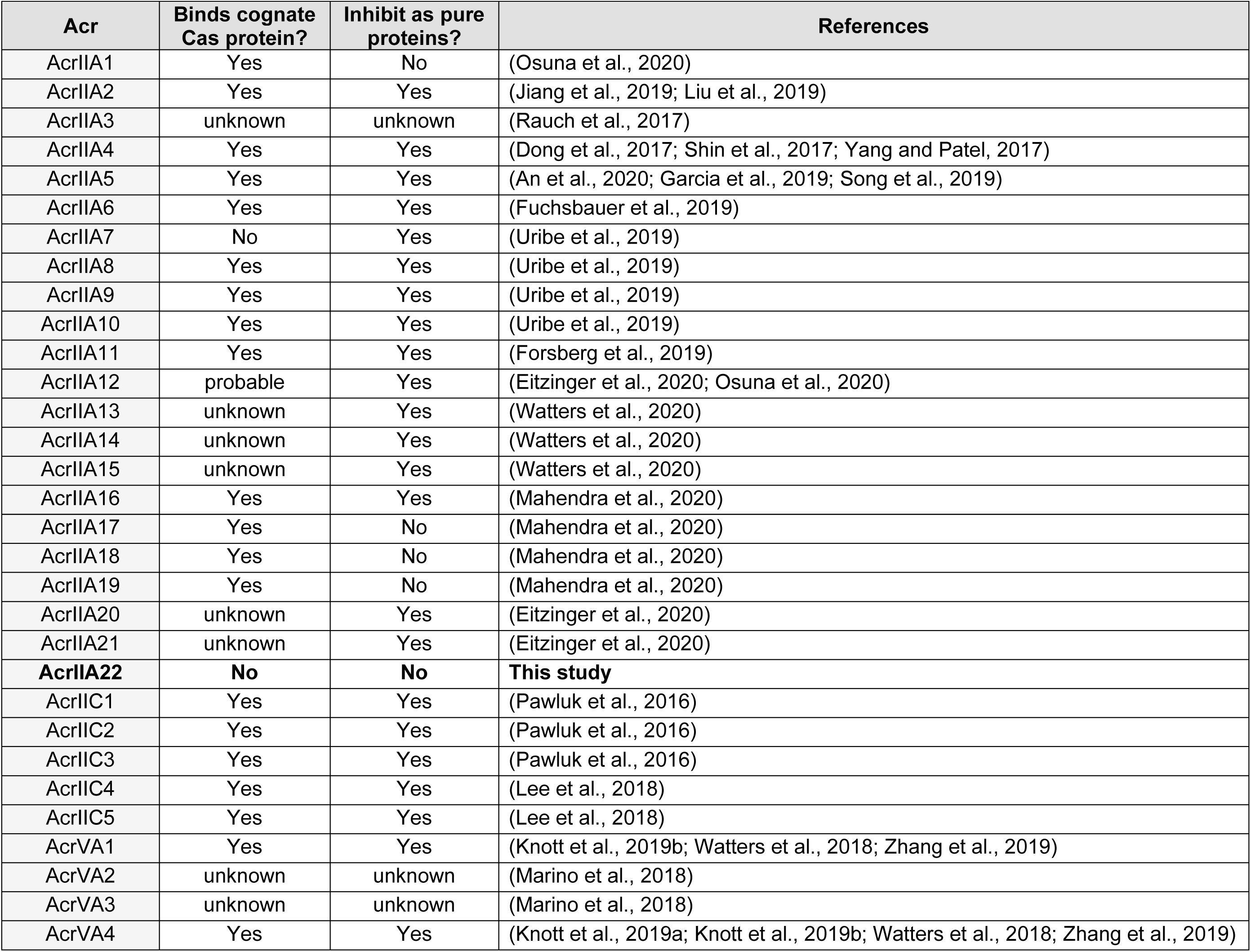
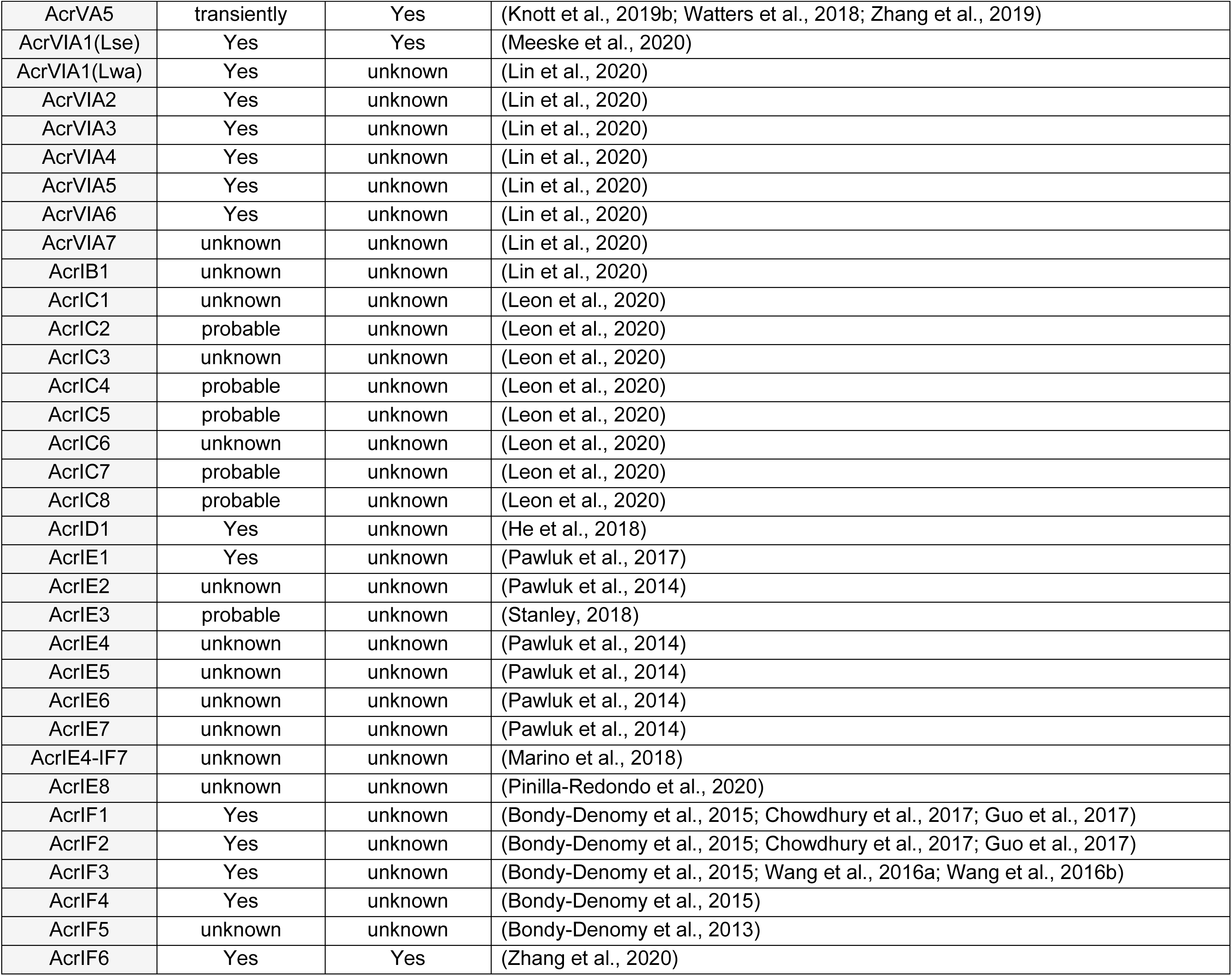
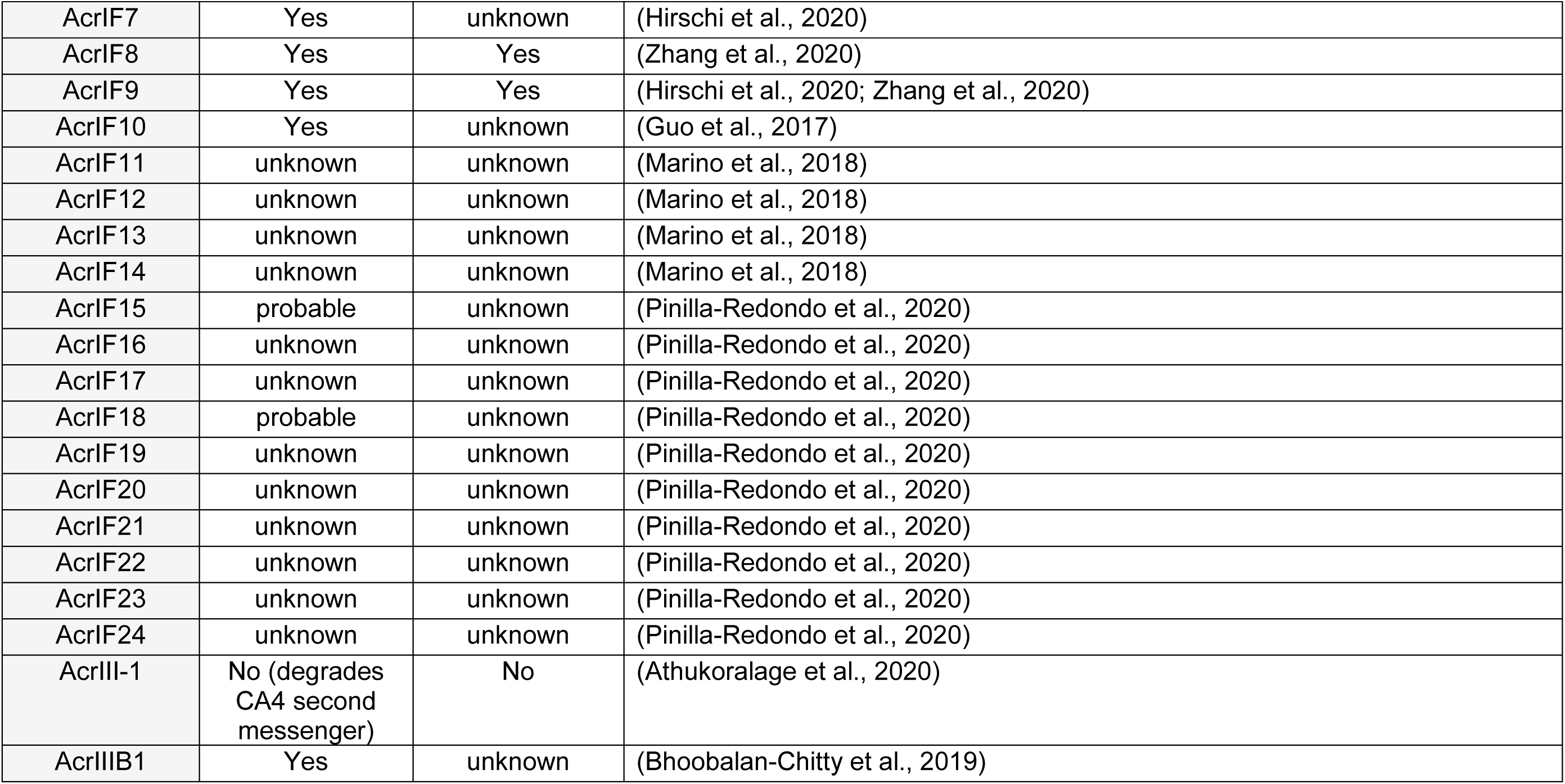
Whether anti-CRISPRs are known to bind Cas proteins and inhibit their cleavage activity as purified proteins.

**Supplemental Table 2.** PC4-like proteins with structural homology to AcrIIA22

| Structural Homolog |  | Function | Similarity to AcrIIA22 |  |  |  |
| --- | --- | --- | --- | --- | --- | --- |
| PDBID | Name | DNA/RNA Binding* | Zscore | r.m.s.d. | n-align | % A.A. ID |
| 4bg7 | PC4 putative transcriptional coactivator p15 | DNA | 6.2 | 2.5 | 54 | 15 |
| 3k44 | D. melanogaster Pur-α | DNA/RNA | 5.9 | 2.6 | 47 | 9 |
| 5fgp | Pur-α repeat I and II from D. melanogaster | DNA/RNA | 5.6 | 2.1 | 48 | 8 |
| 3n8b | Pur-α from B. burgdorferi | DNA/RNA | 5 | 2.8 | 48 | 6 |
| 2gje | Mitochondrial RNA Binding Protein (Trypanosoma brucei) | RNA | 4.9 | 2.5 | 52 | 8 |
| 5zkl | Protein of unknown function SP_0782, S. pneumoniae | DNA | 4.7 | 3.6 | 52 | 12 |
| 5fgo | D. melanogaster Pur-α repeat III | No info | 4.5 | 2.7 | 44 | 14 |
| 1pcf | Replication & transcription cofactor PC4 CTD | DNA | 4.5 | 2.5 | 45 | 7 |
| 2ltt | Putative Uncharacterized Protein YDBC | DNA | 4.5 | 2.8 | 50 | 12 |
| 4bhm | MoSub1-DNA PC4 transcription cofactor | DNA | 3.9 | 2.8 | 45 | 4 |
| 3cm1 | SSGA-like sporulation specific cell division protein | No info | 2.8 | 3.7 | 47 | 13 |
| 1l3a | Transcription factor PBF-2 (P24, WHY1) | DNA | 2.8 | 5 | 48 | 8 |
| 4ntq | Anti-toxin CdiI, E. cloacae | No info | 2.7 | 3 | 49 | 12 |
| 3n1k | WHY2 transcription factor, S. tuberosum | DNA | 2.6 | 2.8 | 52 | 4 |

\*RNA/DNA binding data from (Janowski and Niessing, 2020).

**Supplemental Table 3.**
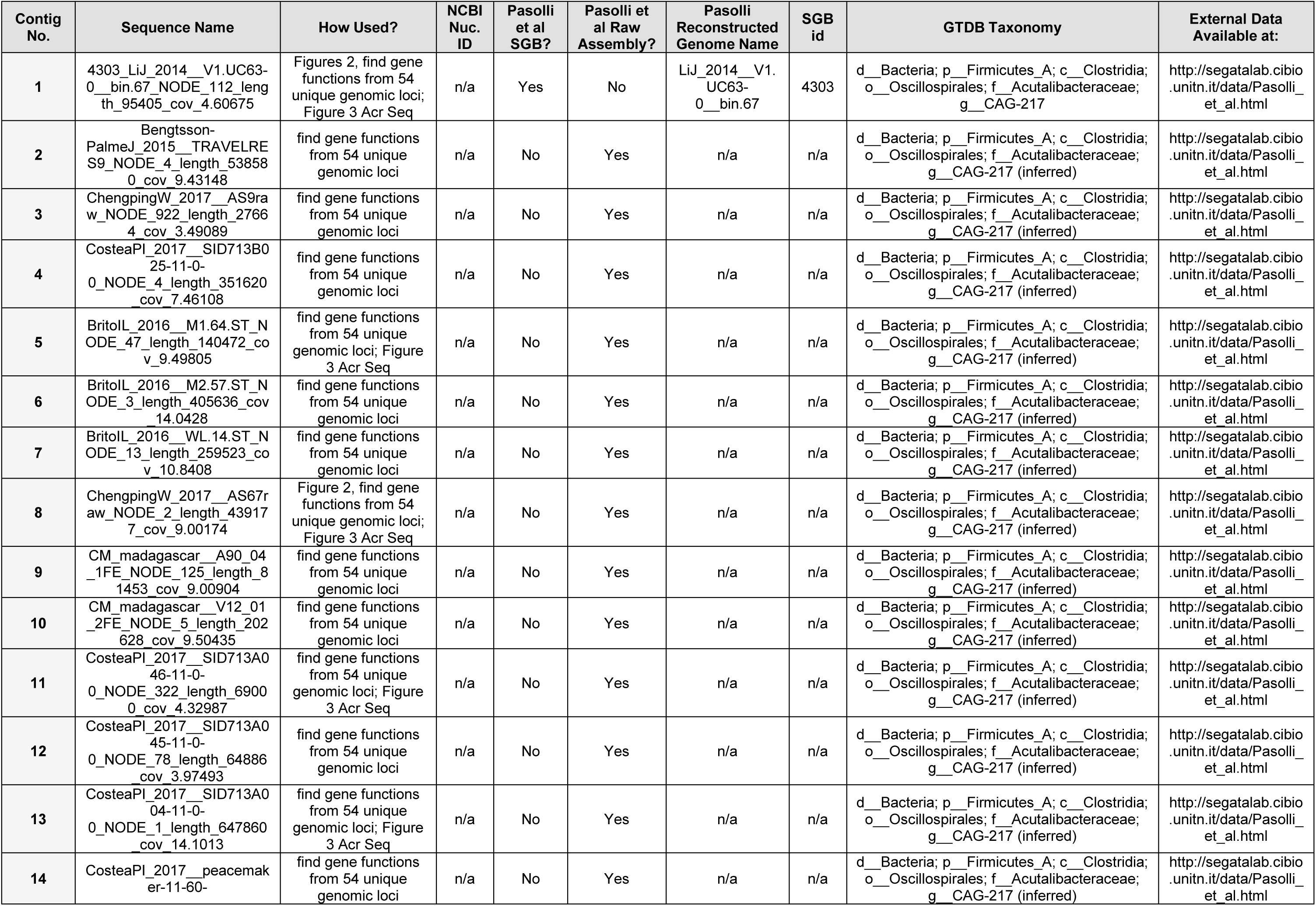
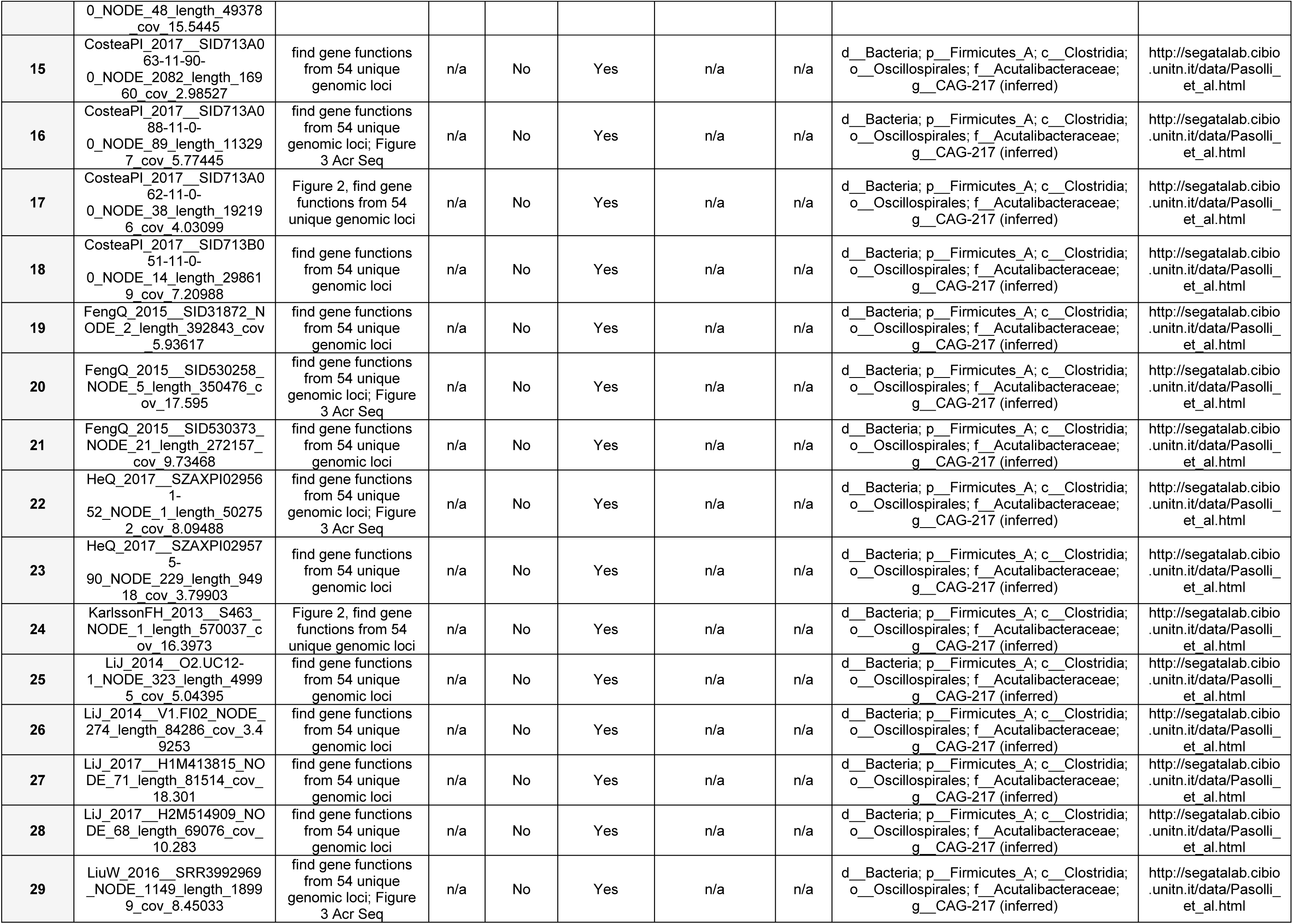
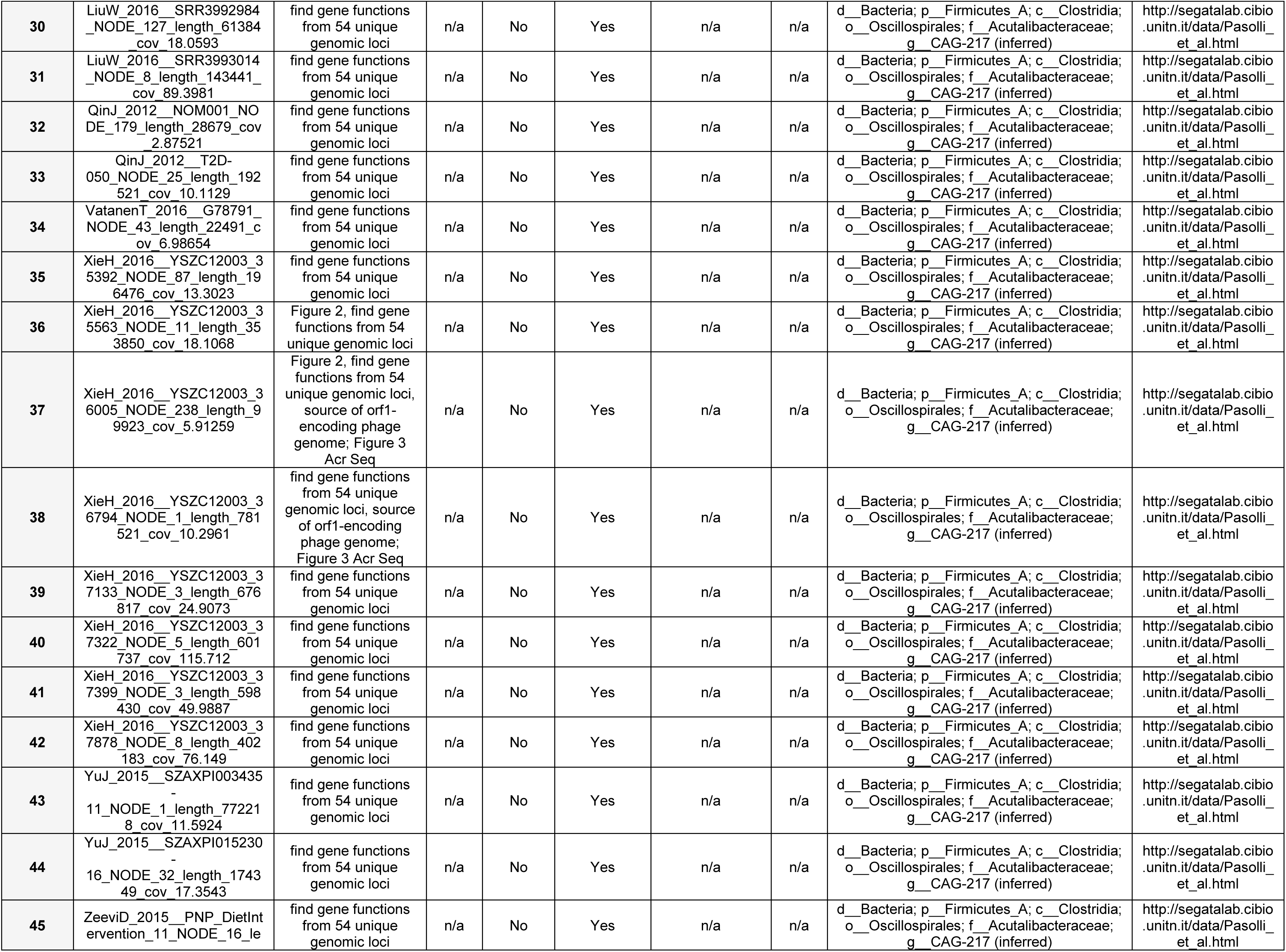
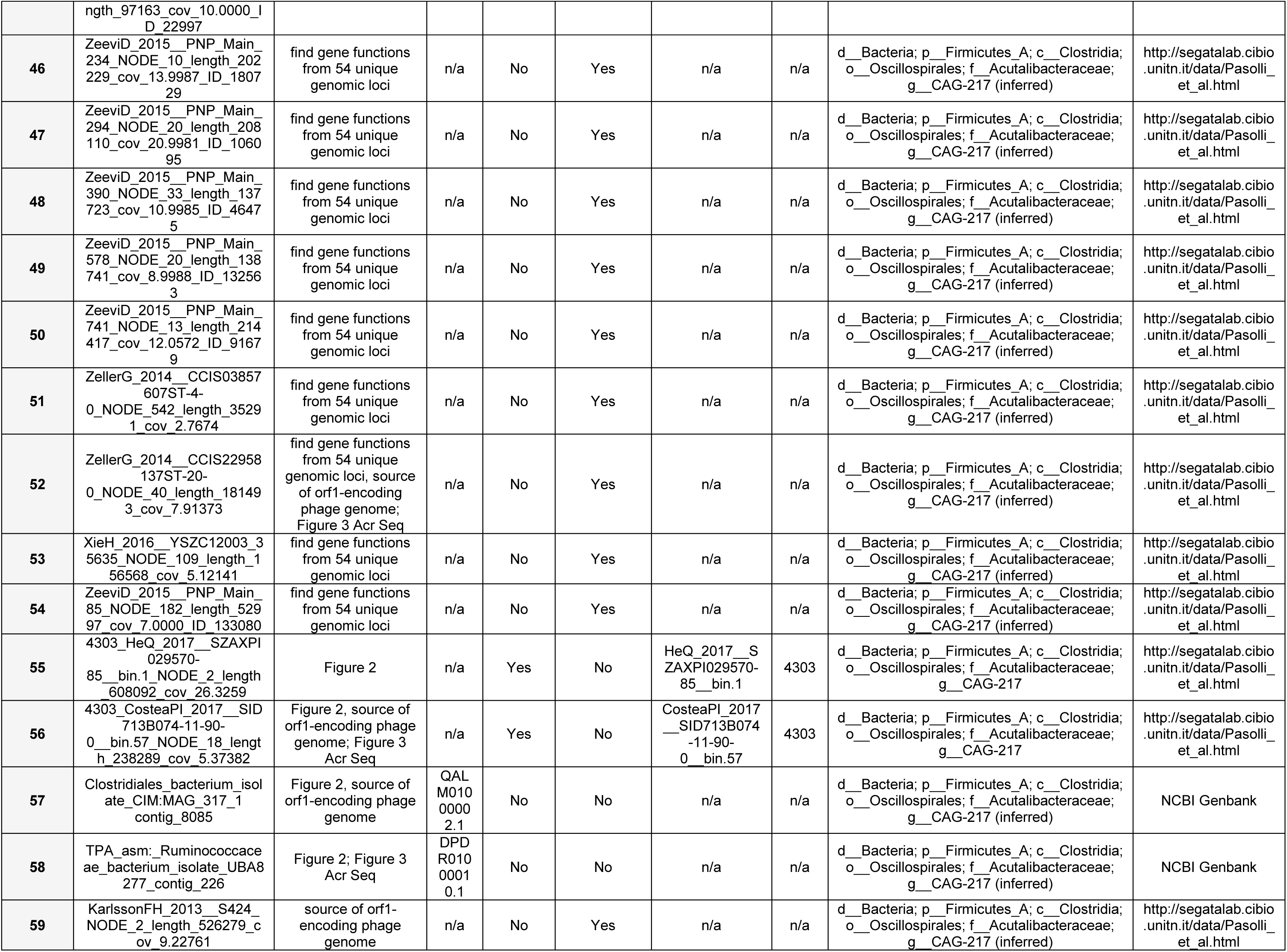
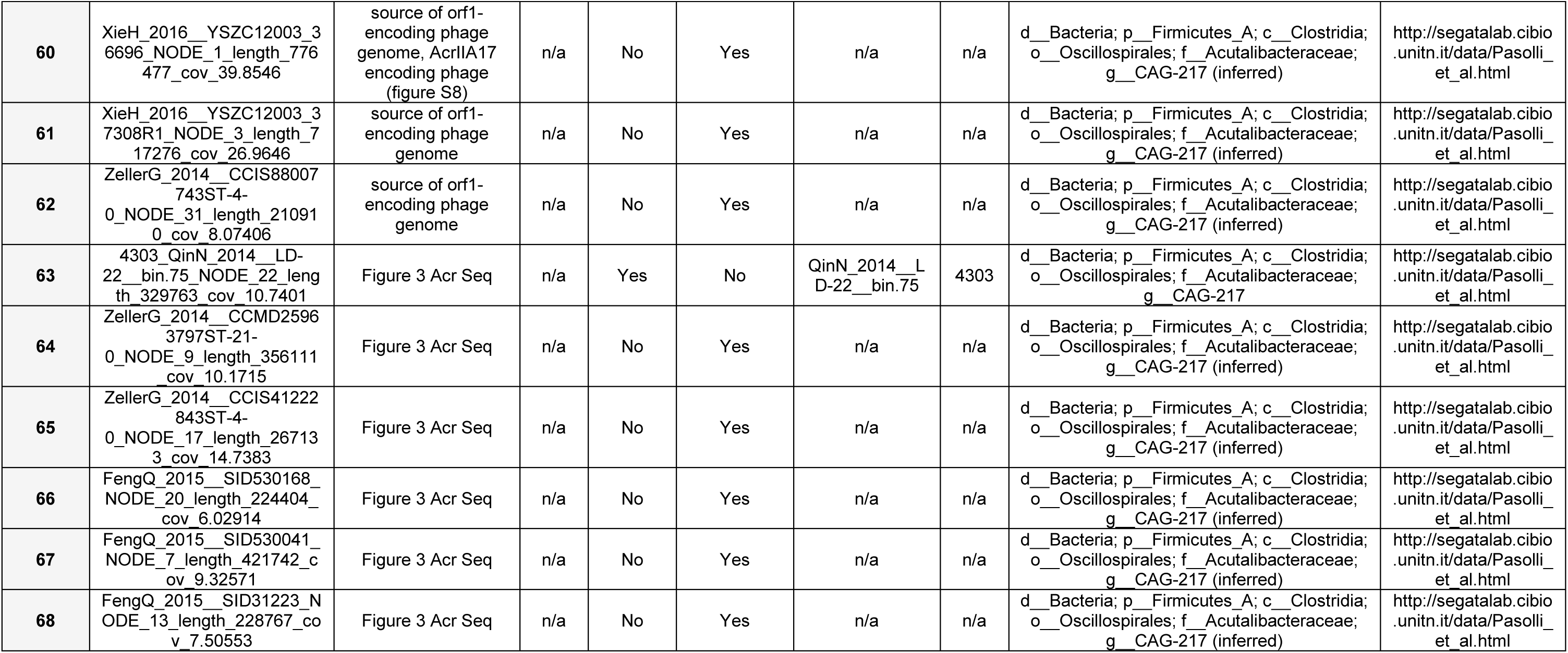
All sequences used in this study. Sequence names and databases are indicated. All sequences and annotations are also available as supplemental data. Sequences retrieved from Pasolli *et al.* refer to the following study: (Pasolli et al., 2019).

**Supplemental Table 4.**
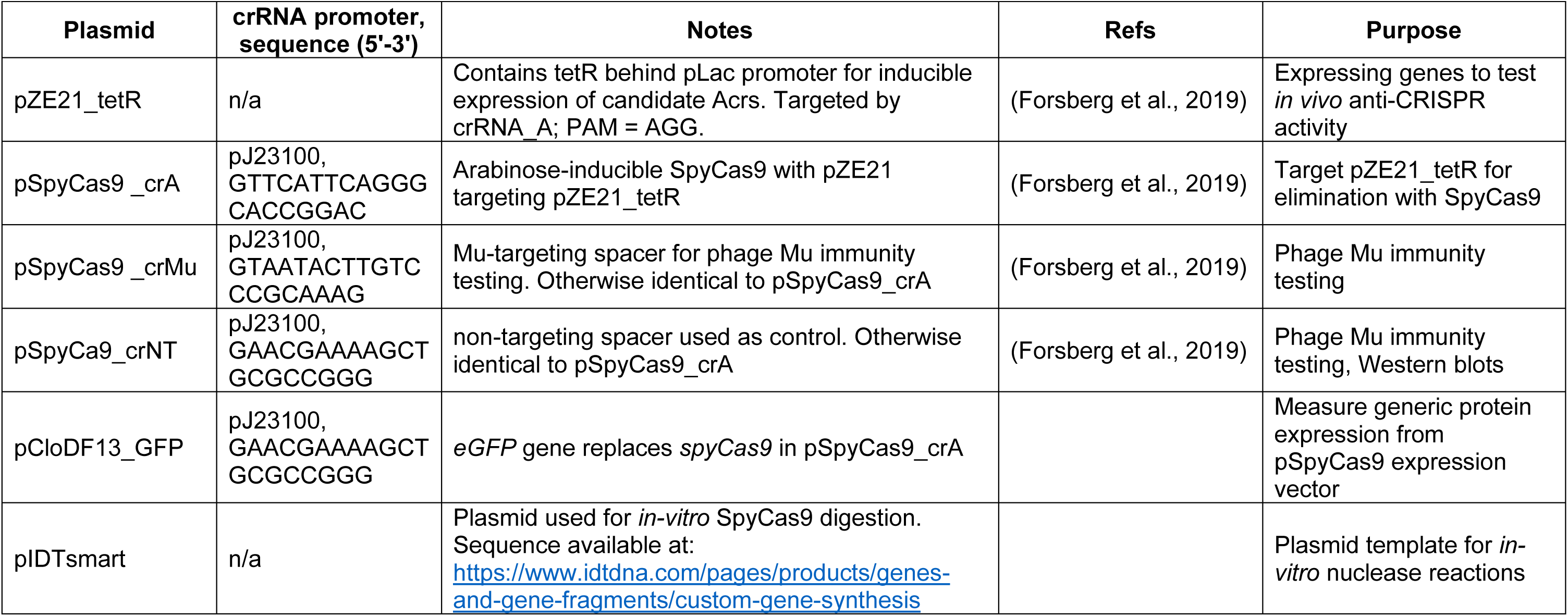
Plasmids used in this study. Supplemental Table S5 indicates genes expressed from pZE21_tetR.

**Supplemental Table 5.**
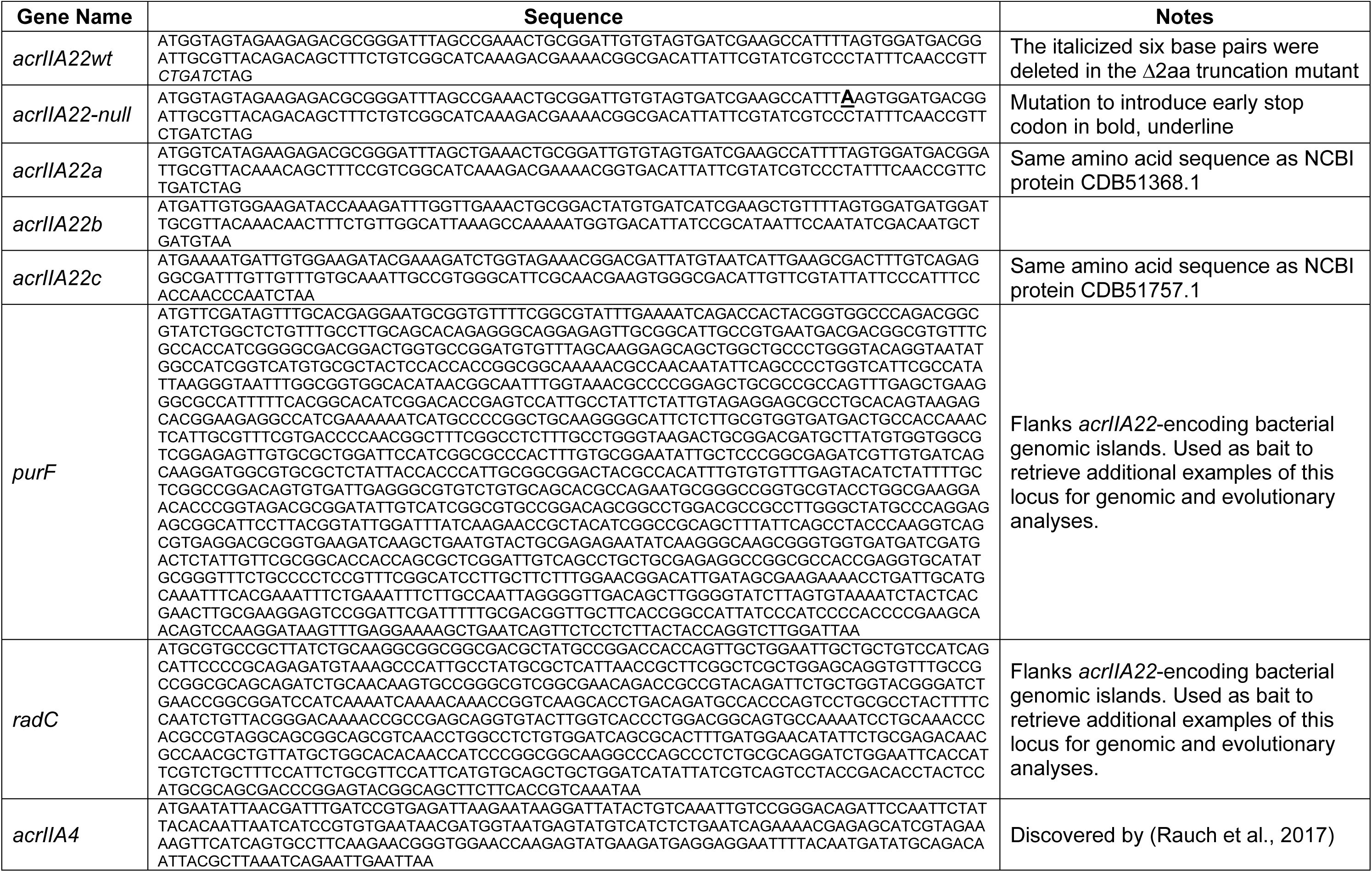
Gene sequences used in this study.

